# SR2: Sparse Representation Learning for Scalable Single-cell RNA Sequencing Data Analysis

**DOI:** 10.1101/2023.07.31.551228

**Authors:** Kai Zhao, Hon-Cheong So, Zhixiang Lin

## Abstract

Single-cell RNA-sequencing (scRNA-seq) technology has been widely used to measure the transcriptome of cells in complex and heterogeneous systems. Integrative analysis of multiple scRNA-seq data can transform our understanding of various aspects of biology at the single-cell level. Many computational methods are proposed for data integration. However, few methods for scRNA-seq data integration explicitly model variation from heterogeneous biological conditions for interpretation. Modeling the variation helps understand the effect of biological conditions on complex biological systems.

Our study proposes SR2 to capture gene expression patterns from heterogeneous biological conditions and discover cell identity simultaneously. Therefore, it can uncover the effect of biological conditions on the gene expression of cells and simultaneously achieve state-of-the-performance in cell identity discovery in our comprehensive comparison. Notably, SR2 is extended to model the effects of biological conditions on gene expression for cell populations, thus uncovering the effect of biological conditions on gene expression for cell populations and identifying putative condition-associated cell populations. To improve its scalability, we incorporate a batch-fitting strategy to ensure it is scalable to scRNA-seq data with arbitrary sample sizes. Moreover, the broad applicability of SR2 in biomedical studies has been demonstrated via applications. The complete package of SR2 is available at https://github.com/kai0511/SR2.

## Introduction

The scRNA-seq technology has emerged as a popular and powerful tool for profiling the transcriptomic landscape of individual cells within complex and heterogeneous systems. It has revolutionized our ability to dissect and understand various aspects of biology at the single-cell level, including developmental biology [1] and gene regulation [2], and opened new avenues for exploring developmental biology, gene regulation, tissue heterogeneity, disease mechanisms, and evolutionary dynamics.

Usually, scRNA-seq studies involve integrative analysis of transcriptomic data of individual cells measured with different technologies and derived from tissues [1] of individuals with different phenotypes [2]-[4] across different conditions [5] and even species. Indeed, the integrative analysis of heterogeneous scRNA-seq datasets to identify cell types or states and to compare gene expression across biological conditions has tremendous potential to transform our understanding of complex and heterogeneous biological systems [6]. A successful example of this practice is that joint analysis of scRNA-seq data from multiple melanoma tumors identifies an immune resistance program in malignant cells, which predicts clinical responses to immunotherapy in melanoma patients [7]. However, the technical variation from different sequencing platforms or library-preparation protocols and the biological heterogeneity inherent in biosamples under different biological conditions, such as donors, tissues, or phenotypes, make the integrative analysis of single-cell RNA-seq datasets challenging. Conceptually, the technical variation and biological heterogeneity across the scRNA-seq dataset are considered batch effects.

To address these challenges in the integrative analysis of scRNA-seq data, computational methods or statistical approaches have been developed and differ in their implications for analysis. On the one hand, computational approaches, including BBKNN [8], FIRM [9], and FastMNN[10], are proposed, which assume that the batch effect in scRNA-seq data is almost orthogonal to the biological subspace [10]. Thus, their ability to correct batch effects originating biologically across scRNA-seq datasets is limited [11]. On the other hand, other computational methods [6], [11]-[16] have been developed to address the biologically originated batch effects. Seurat [17], probably the most popular among them, matches cell states across biosamples by a shared correlation space defined by canonical correlation analysis. Liger [6] employs nonnegative matrix factorization (NMF) to delineate shared and dataset-specific features of cells from different single-cell samples. Harmony [14] integrates single-cell data by projecting cells into a shared embedding. Scanorama [18] leverage matches of cells with similar transcriptional profiles across scRNA-seq datasets to perform batch correction and integration. The scINSIGHT [11] employs additive NMF to model the variations from heterogeneous biological conditions to perform scRNA-seq data integration.

However, most of these computational methods for single-cell data integration concentrate on technical variation harmonization and cell identity annotation. Few of them explicitly account for the variation arising from heterogeneous biological conditions. As a result, they cannot provide direct model interpretation on the effect of biological conditions on gene expression over cell populations. To achieve the aim, scINSIGHT [11], based on NMF, probably is the first study modeling the variations from heterogeneous biological conditions in data integration, but its scalability is unsatisfactory as it cannot yield results in two days when applied to scRNA-seq data with sample size ∼23000. With the availability of massive scRNA-seq datasets, the scalability issue of scINSIGHT will be more prominent. Moreover, the ability of NMF to directly infer down-regulations of biological elements in biological data is unsatisfactory [19]. In fact, different biological conditions exert different effects on the gene expression of cells and cellular states of different cell populations (types). To our knowledge, very few or no computational tools for scRNA data analyses directly model the effect of biological conditions on different cell populations in low-rank latent spaces for interpretation. Explicitly modeling the condition-specific variation across different cell populations enables interpretable analysis of the integrated data, such as revealing the effect of biological conditions on gene expression for cell populations and identifying condition-associated cell populations, thus gaining insights into the effects of biological conditions on complex and heterogeneous biological systems.

To fill these gaps, we propose a novel statistical framework, SR2, based on an ensemble of matrix factorization and sparse representation learning. In SR2, we leverage matrix factorization to capture variation from different biological conditions (e.g., donor, tissue, and disease status) within a shared low-rank latent space and sparse representation learning to decompose cellular variations across biosamples simultaneously. The rationale behind our modeling is that different donors demonstrate heterogeneity in gene expression via pathways and that biological conditions can affect the gene expression of the donors via the same pathways. After modeling the heterogenous biological variations, sparse representation learning is incorporated to obtain low-rank embeddings for cells across different biosamples in another shared low-rank latent space. Then, the technical artifacts can be controlled because the artifacts are usually not shared across biosamples. To ensure scalability and efficiency, SR2 incorporates *a batch strategy* in model fitting, which improves the speed of computation and reduces memory consumption. This allows for analyzing large-scale scRNA-seq datasets without sacrificing performance. Furthermore, we extend SR2 to explicitly model the effect of biological conditions on different cell populations (types). Through this extension, SR2 reveals how different biological conditions influence gene expression among different cell populations, thus identifying *condition*-associated cell populations. Details of our method are presented in the Method section.

In contrast to existing computational methods, SR2 has the following virtues. Firstly, it is a general and flexible framework for integrative analysis of scRNA-seq datasets, considering heterogeneous biological variation from *multiple* biological conditions. In SR2, biological conditions can be defined based on research questions. Secondly, the extension of SR2 proposed to model the effect of biological conditions on gene expression among different cell populations (types), thus gaining insight into the heterogeneous effect of biological conditions on cell populations and identifying *condition-associated* cell populations. Thirdly, modeling variation from different biological conditions over or across cell populations provides informative inputs for interpretable downstream analyses, such as linking gene activities to biological conditions, revealing biological processes (BPs) underlying biological conditions, uncovering the molecular characteristics of donors, revealing the effects of biological conditions on gene expression for different cell populations, and identifying *condition-associated* cell populations. Fourthly, it offers two different fitting strategies tailored to analyze scRNA-seq data with a wide range of sample sizes. The whole data fitting strategy allows efficient analysis of scRNA-seq datasets with small or median sample sizes, and the batch strategy ensures it is scalable to scRNA-seq datasets with *arbitrary* sample sizes. Moreover, it allows missing values in the gene expression matrix, which dramatically improves the applicability of SR2. Finally, it achieves state-of-the-art performance in identifying cell populations across biosamples. Its wide applicability is demonstrated via intensive applications to various biomedical scenarios. Further extension of SR2 includes large-scale analysis of other single-cell omics data.

## Results

### Overview of SR2

We propose a flexible computational method SR2 (sparse representation learning for scalable single-cell <u>R</u>NA sequencing data analysis), presented in Figure 1. Here we use donor and phenotype as an example of biological conditions for illustration. In scRNA-seq studies, the RNA expression of cells from different donors with different phenotypes is profiled (Figure 1A). To facilitate interpretable scRNA-seq data analysis, SR2 models variation from multiple biological conditions (e.g., donor and phenotype) with matrix factorization and cellular variation with sparse representation learning (Figure 1B).

**Figure 1.**
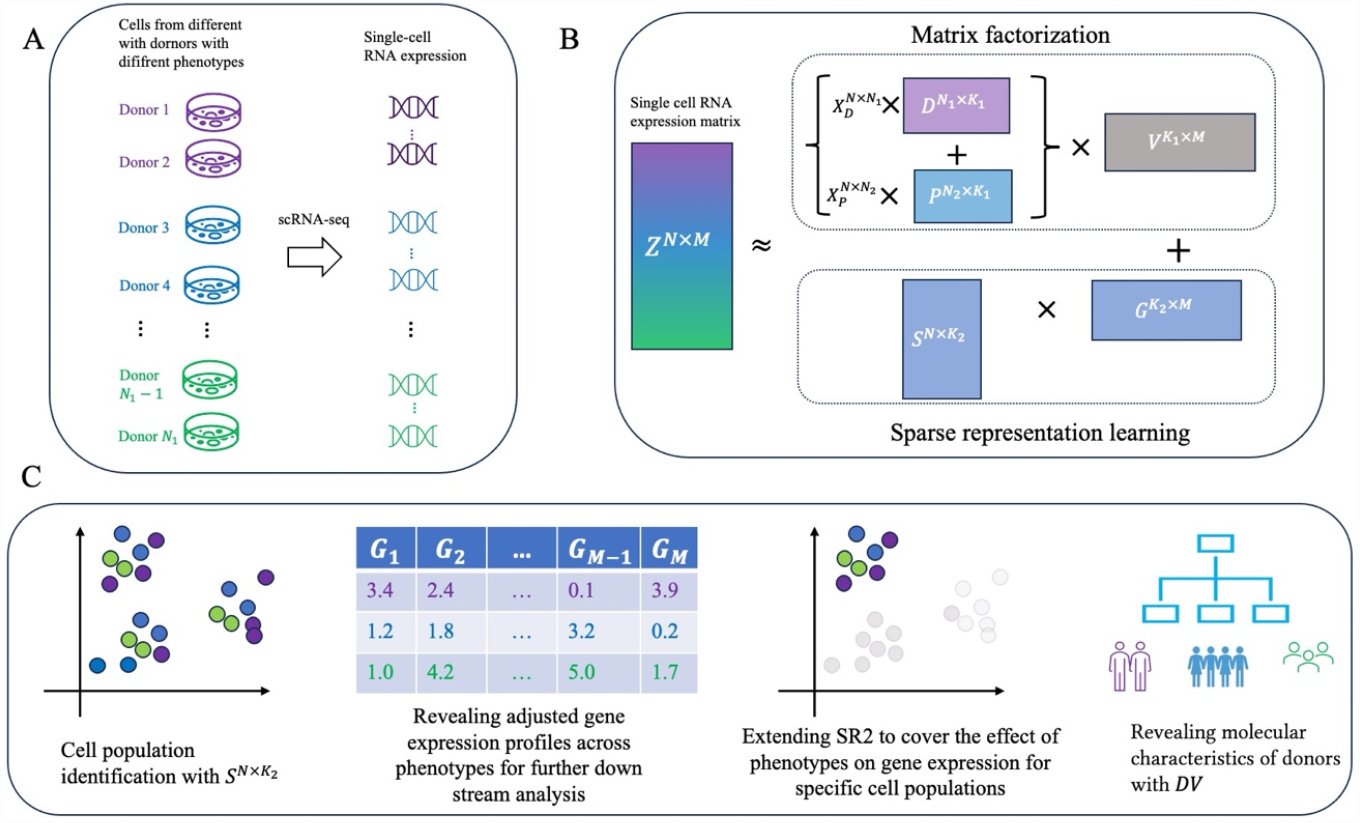
Overview of SR2 A. The scRNA-seq data is obtained from biosamples from multiple biological conditions. Variations from biological conditions (e.g., donor and phenotypes) and technical artifacts bring batch effects in the scRNA-seq data matrix. B. SR2 models variation from donor and phenotype with matrix factorization and cellular variation with sparse representation learning. C. The output from SR2 enables various downstream analyses, such as cell population annotation, revealing the effect of phenotypes on gene expression over all cell populations, and uncovering donors’ molecular characteristics. SR2 can also be extended to reveal the effect of biological conditions (e.g., phenotypes) on gene expression for different cell populations.

In SR2, the expression level of gene *m* for sample *i*, which is obtained from donor *j* with phenotype *t, z*_*im*_, can be modeled as

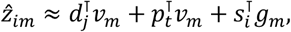

where *d*_*j*_, *p*_*t*_, *ν*_*m*_ are vectors of length *K*_1_, *s*_*i*_, *g*_*m*_ are vectors of length *K*_2_. The donor and phenotype information of samples is available. The equation has the following matrix representation

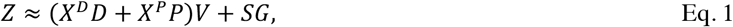

where 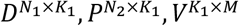 are matrices of latent representations for *N*_1_ donors, *N*_2_ phenotypes, and *M* genes, respectively, 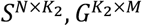 denote latent representations for *N* cells and *M* genes after modeling variation from donor and phenotype, and *X*^*D*^, *X*^*P*^ are indicator matrices of *N* rows and *N*_1_ and *N*_2_ columns, respectively, which are the dummy variables for *N* samples. The objective function for the equation with matrix representation is as follows:

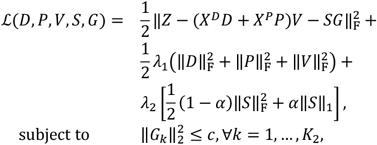

where *G*_*k*_ is the *k*-th row of matrix *G. c* is a constant, restricting the scale of each component of *G, λ*_1_, *λ*_2_, *α* are tuning parameters, and ‖⋅‖_*F*_ represents the Frobenius norm. Alternating block coordinate descent (BCD) is utilized to optimize the objective. The technical details of the optimization of SR2 are presented in the Methods section.

The output from SR2 provides informative input for various downstream analyses. First, cell populations (types) can be identified with *S* from Eq. 1, the effect of phenotypes on gene expression over cell populations can be revealed by multiplying *P* with *V*, and the molecular characteristics of donors can be uncovered by examining donor representation *D*. More importantly, SR2 can be extended to reveal the effects of biological conditions (e.g., phenotypes) on gene expression for different cell populations, thus identifying phenotype-associated cell populations via functional analysis. The demonstrations of these analyses are presented below.

### Datasets

In this study, we first applied SR2 with the whole data strategy to three scRNA-seq data from studies on the pancreas of diabetic and normal donors (DM dataset) [2], human airway epithelium of donors with different smoking habits (Smoking dataset) [5], and peripheral blood cells of patients experiencing mild to severe COVID-19 infection (COVID-19 dataset) [3]. The sample size of three datasets ranges from ∼6000 to over 30000. Moreover, to empirically demonstrate the scalability and performance of SR2 in analyzing large-scale scRNA-seq data, we further applied SR2 with the batch strategy to the immune dataset, which offers scRNA-seq data containing more than 300,000 immune cells extracted from 16 different tissues of 12 deceased adult organ donors [1], and the GBM (glioblastoma) data of over 200, 000 human glioma, immune, and other stromal cells from glioma patients [4].

In the above analyses, cell populations for each dataset were identified. Details on cell population identification are provided below. Hence, we investigated the effects of biological conditions on gene expression for cell populations with the extension of SR2 presented in the Method section. Details of the analysis are presented below.

### SR2 links the activities of genes to biological conditions

Currently, most computational methods for single-cell analysis focus on cell type annotation, and hence they cannot uncover gene expression levels across biological conditions. However, SR2 captures variation from heterogeneous biological conditions in scRNA-seq data. This practice enables linking gene activities with biological conditions in analyzing scRNA-seq data.

To illustrate with Eq. 1, the adjusted expression profiles of the *M* genes for phenotypes can be obtained by multiplying *P* by *V*, which reveal gene expression changes across different phenotypes. Note variation from donors is controlled in the above processes as we consider donor variation in SR2. Moreover, phenotype-associated genes can be identified by examining the correlation between phenotype representations and gene representations. This idea is inspired by one previous study [20]. Practically, *p*-values in the calculation are adjusted for multiple statistical testing.

For demonstration purposes, we carried the analysis to the DM dataset, COVID-19 dataset, and GBM dataset and visualized expression changes of the most variable genes across phenotypes (Figures 2A, 2B, and 2C, respectively) and the top phenotype-associated genes (Figures 2D, 2E, and 2F, respectively) for the three datasets.

**Figure 2.**
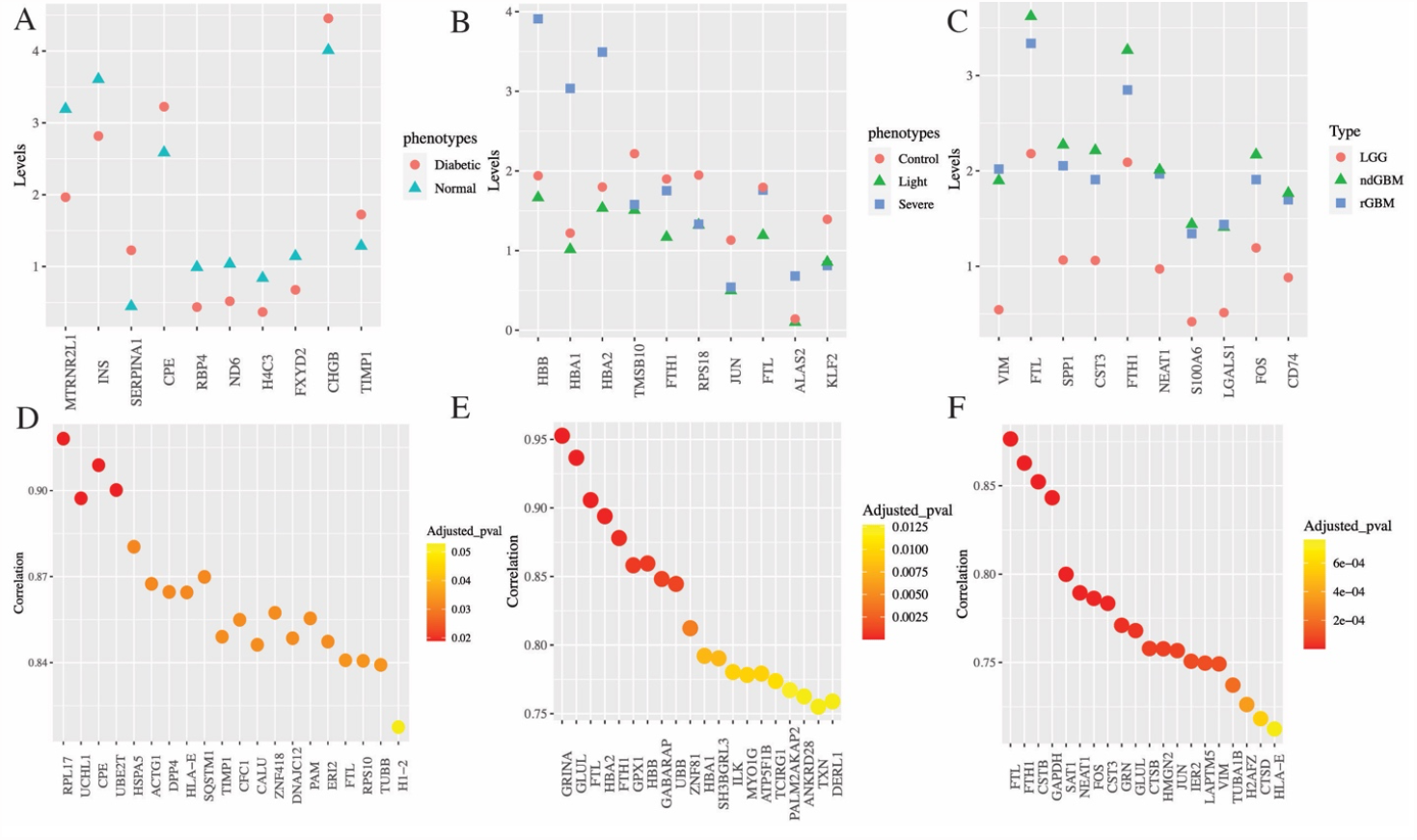
SR2 links activities of genes to phenotypes of donors A. Changes in the expression of the most variable (top 10) genes across diabetic and normal conditions from analysis of the DM dataset are shown. B. Changes in the expression of the most variable (top 10) genes across control, light, and severe COVID-19 from analysis of the COVID-19 dataset are shown. C. Changes in the expression of the most variable (top 10) genes across different types of GBM from analysis of the GBM dataset are shown. The words “LGG”, “ndGBM”, and “rGBM” stand for low-grade gliomas, newly diagnosed GBM, and recurrent GBM, respectively. D. The top 20 diabetes-associated genes ranked by adjusted *p*-values are presented. E. The top 20 severe COVID-19-associated genes ranked by adjusted *p*-values are displayed. F. The top 20 recurrent GBM-associated genes ranked by adjusted *p*-values are presented.

The expression levels of the ten most variable genes across diabetic and normal conditions are presented in Figure 1A. The relationship between the top genes from Figure 1A, such as MTRNR2L1 [21], INS [22], SERPINE1 [23], and CPE [24], and diabetes have been studied previously. Moreover, top statistically diabetes-associated genes (Figure 1D), such as RPL17, differentially expressed between T2DM and normal samples [25], UCHL1, exacerbating human islet amyloid polypeptide toxicity in β-cells [26], and CPE [24] have also been studied.

Regarding severe COVID-19, HBB (Figures 2B and 2E) is identified by a previous study [24] in differential expression analysis of severe COVID-19 against normal and COVID-19 with scRNA-seq data. Moreover, expression levels of HBA1 (Figures 2B and 2E) and HBA2 (Figure 2B), which produce alpha globin carrying oxygen to cells and tissues throughout the body, are high in severe COVID-19. Another two genes, GRINA and FTL (Figure 2E), were also differentially expressed between severe COVID-19 patients and controls [27], [28].

For our analysis of GBM, there is a wide difference in gene expression levels between GBM and LGG, compared with the difference between recurrent GBM and newly diagnosed GBM (Figure 2C). Also, genes listed in Figure 2C and/or Figure 2F are shown to be associated with GBM, such as FTL [29], FTH1[30], GAPDH [31], and VIM [32].

### SR2 reveals the gene expression patterns for cell populations across biological conditions

Previously, we presented the expression levels of genes under various biological conditions. However, it is important to note that gene expression for different cell populations can be influenced differently by the same biological condition. Therefore, we utilized the extended SR2 to uncover the heterogeneous effects of biological conditions on gene expression for different cell populations.

For illustration, assume that there are *N*_*c*_ cell populations (or types) and *N*_2_ levels for the biological condition of interest. Let *X*^*c*^ denote the matrix of dummy variables for all combinations of biological conditions and cell populations (or types). Mathematically, we consider 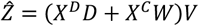 to achieve our aim. In the equation, *X*^*D*^, *D* denote the indicator matrix of dummy variables for donors and donor representation, respectively, and 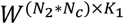 is low-rank representations of cell populations under different biological conditions, while controlling variation from donors. After optimizing the defined problem, we can multiply *W* with *V* to obtain adjusted expression profiles for cell populations under different biological conditions. Then, expression levels for specific cell populations across biological conditions can be revealed. Moreover, functions of one cell population under a specific biological condition can also be identified by examining the biological processes (BPs) enriched by genes in the upper or lower 5% quantile of the adjusted expression profile for the cell population under the biological condition.

To implement the above concept, we applied the extended SR2 approach to analyze the DM and COVID-19 datasets and then multiplied *W* with *V* to reveal expression profiles for cell populations across biological conditions. Subsequently, change in expression levels of the top 10 variable genes for the selected cell population across biological conditions are depicted in Figure 3. In our analysis of the two datasets, we utilized the cell types provided by the original dataset to annotate the cell populations identified by low-rank embeddings from SR2. Details on cell population annotation are presented in the subsequent section. Moreover, the proportion and cell type of cell populations for the DM and COVID-19 dataset is provided in Supplementary Tables S1 and S2, respectively.

**Figure 3.**
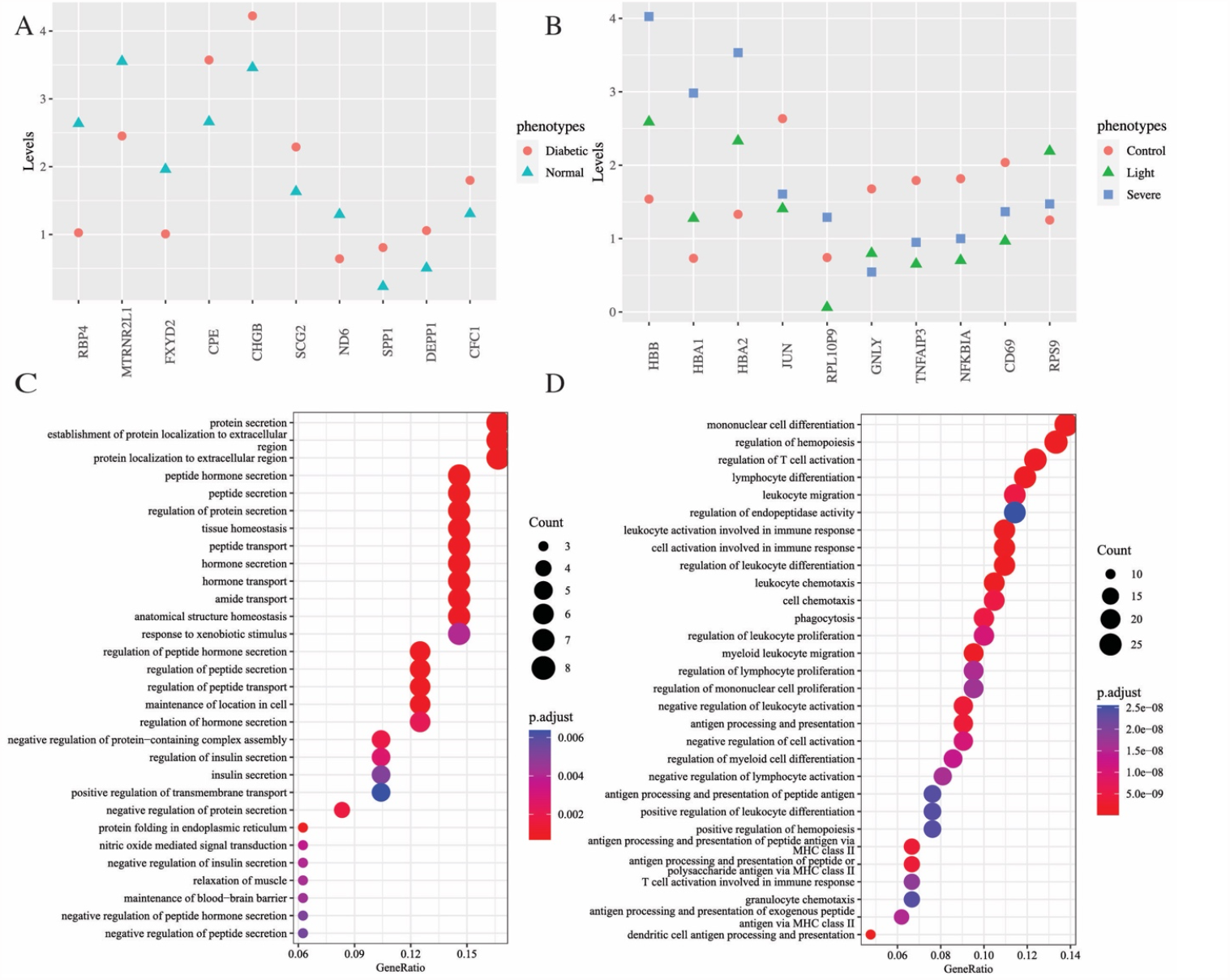
SR2 reveals expression changes in the top variable genes for cell populations across phenotypes and the function of cell populations A. The figure shows changes in the expression levels of the most variable (top 10) genes for the largest beta cell population across control and diabetes with the DM dataset. B. The figure shows changes in the expression of the most variable (top 10) genes for the T cell population across control, light COVID-19, and severe COVID-19 with the COVID-19 dataset. C. The top 30 BPs enriched by top genes in the adjusted expression profiles for the largest beta cell population from normal donors are displayed. D. The top 30 BPs enriched by top genes in the adjusted expression profiles for the T cell population from donors with severe COVID-19 are displayed.

Regarding the analysis of the DM dataset, three beta cell populations were identified (Table S1). Interestingly, the proportion for two larger beta cell populations is lower in diabetic donors than normal donors, but the proportion of the other (smallest) beta cell population is higher in diabetic donors than in controls. First, we examined changes in the expression of the most variable genes (top 10) for the largest beta cell population (Beta cell population 1 in Table S1) across control and diabetes (Figure 3A). We observed noticeable changes in the expression levels of MTRNR2L1 and CPE in the beta cell population across diabetes and normal (Figure 3A) and rediscovered some genes identified in Figure 2A previously. Moreover, the importance of the identified genes, such as RBP4 [33], FXYD2 [34], [35], and CHGB [36], in beta cells and their involvement in diabetes have been studied previously. These findings suggest that the gene expression difference in the beta cell population across phenotypes contributes to the overall gene expression difference across phenotypes, implying the contributing role of beta cells in diabetes. Subsequently, in the enrichment analysis of the three beta cell populations across diabetes and controls, top genes from the two larger ones were found to be enriched for insulin secretion and regulation (Figure 3C and S2A), but the smaller beta cell population did not exhibit these insulin-related BPs (Figure S2B). Together with the distribution of the three beta populations (Table S1), our findings imply that diabetes is associated with a decrease in the populations of beta cells that secrete insulin and an increase in the population of beta cells that do not participate in insulin secretion. This further emphasizes the role of beta cells in diabetes. In the work, we will further clarify the role of beta cells in diabetes.

In the analysis of the COVID-19 dataset, we noticed that the proportion of T cells is significantly lower in donors with severe COVID-19 than those in mild COVID-19 or controls (Table S2). To delve deeper into the characteristics of T cells in COVID-19, we investigated the expression changes of the most variable genes for the T cell population across different COVID-19 conditions (Figure 3B). The relationship between HBB, HBA1, and HBA2 (Figure 3B) and COVID-19 severity has been discussed in one previous section. Moreover, two other genes, TNFAIP3 [37] and GNLY [38] (Figure 3B), in T cells have been reported to play an important role in COVID-19. Additionally, the BPs enriched by top genes from T cells of donors with severe COVID-19 involve human immune responses (Figure 3D). These findings highlight the important role of T cells in the human immune response to COVID-19.

### SR2 reveals BPs underlying biological conditions

In one previous section, we revealed how SR2 helps link the activities of genes to biological conditions. A natural question is whether these genes are enriched for BPs to exert effects. Let us illustrate with results from our analysis with the DM dataset. Following the practice from the previous section, we can reveal BPs enriched by down- regulated genes with the largest negative differences in the adjusted expression profiles between T2DM and normal after computing the adjusted expression profiles for diabetes and control. In practice, we utilized the genes in the lower 5% quantile of the difference to obtain down-regulated BPs; likewise, genes in the upper 5% quantile can be used to discover up-regulated BPs.

For demonstration purposes, we revealed down-regulated BPs for cells from donors with T2DM versus those from normal control (Figure 4A), up-regulated BPs for mild COVID-19 versus normal (Figure 4B), and severe COVID- 19 versus mild COVID-19 (Figure 4C), up-regulated BPs for light smoking versus non-smoking (Figure 4D) and heavy smoking versus light smoking (Figure 4E), and up-regulated BPs for recurrent GBM smoking (Figure 4F) and recurrent GBM versus new diagnosed GBM (Figure 4H).

**Figure 4.**
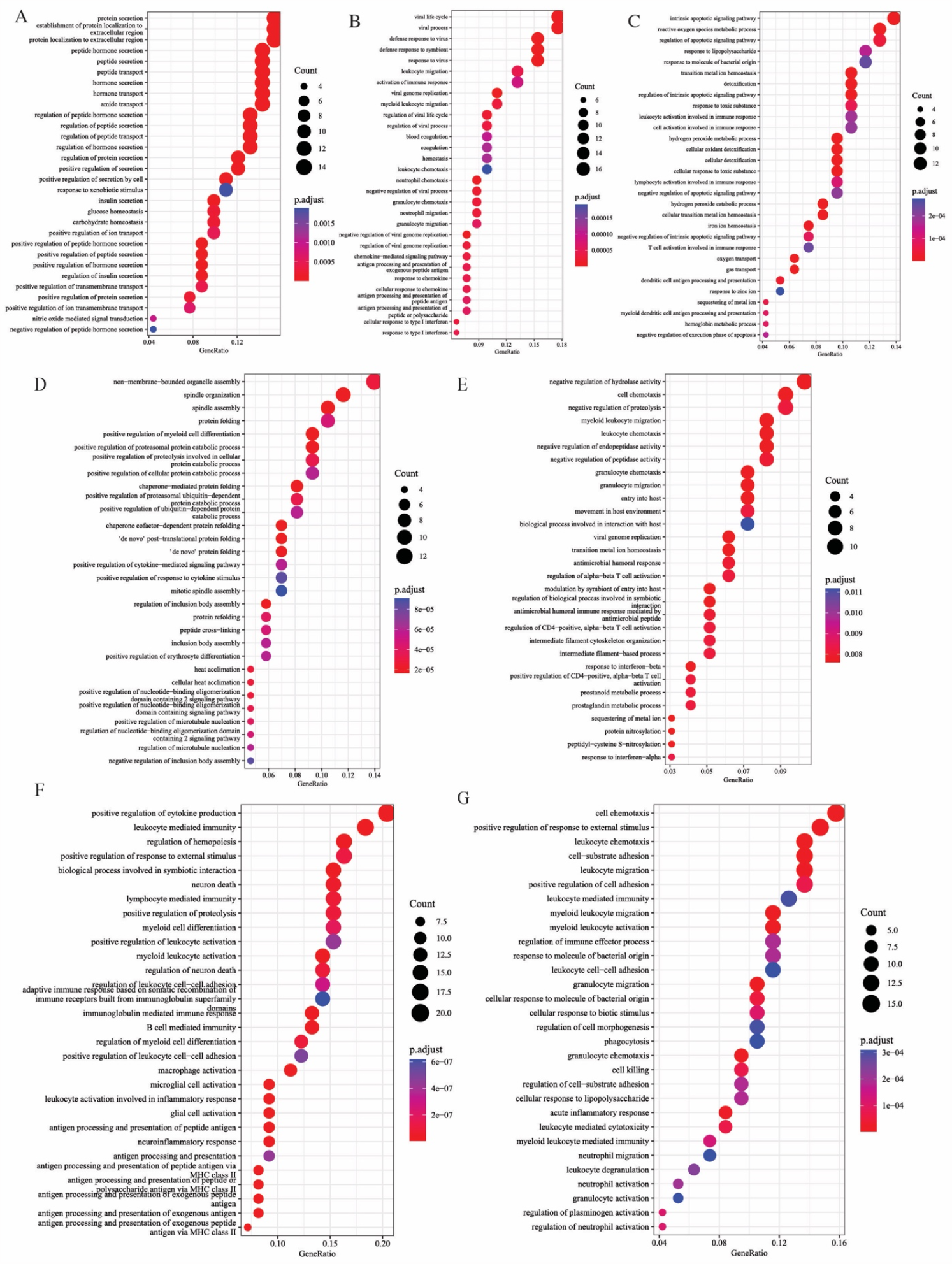
SR2 reveals BPs underlying biological conditions in applications of SR2 A. The down-regulated BPs enriched by genes in the lower 5% quantile of the difference in the adjusted expression profiles between the diabetic and normal conditions are shown. B. The up-regulated BPs enriched by genes in the upper 5% quantile of the differences in the adjusted expression profiles between mild COVID-19 and normal are shown. C. The up-regulated BPs enriched by genes in the upper 5% quantile of the differences in the adjusted expression profiles between severe and mild COVID-19 are shown. D. The up-regulated BPs enriched by genes in the upper 5% quantile of the difference in the adjusted expression profiles between light smokers and non-smokers are shown. E. The up-regulated BPs enriched by genes in the upper 5% quantile of the difference in the adjusted expression profiles between heavy and light smokers are shown. F. The up-regulated BPs enriched by genes in the upper 5% quantile of the adjusted expression for the recurrent GBM are shown. G. The up-regulated BPs enriched by genes in the upper 5% quantile of the difference in the adjusted expression profiles between the recurrent GBM and newly diagnosed GBM are shown.

Regarding T2DM, we rediscovered known diabetes-associated BPs, such as insulin secretion, glucose homeostasis, and carbohydrate homeostasis (Figure 4A). Some other identified BPs involved in peptide secretion, peptide hormone secretion, and amide transport (Figure 4A) are suggested to play important roles in T2DM by previous studies [39], [40]. Moreover, the peptidergic system is received wide attention for its clinical potential to treat obesity [39].

In the analysis of the COVID-19 dataset, up-regulated BPs in cells from donors with mild COVID-19 versus those from normal involve the viral life cycle, viral process, and (defense) response to viral (Figure 4B). Moreover, the BPs related to immune response, such as (myeloid) leukocyte and neutrophil chemotaxis, were also enriched (Figure 4B). Regarding severe COVID-19, we found that BPs related to apoptotic signaling pathways are activated, compared with mild COVID-19 (Figure 4C), consistent with the previous finding that suggests that T cell apoptosis characterizes severe COVID-19 disease [41]. Meanwhile, the BPs involved in detoxification and T-cell activation are also identified (Figure 4C).

Regarding the effects of different smoking behavior, our results uncover the immune activation in cells from heavy smokers compared to those from light smokers (Figure 4E). Moreover, up-regulated BPs involved in the mitotic spindle and protein folding were identified when comparing light smoking to non-smoking (Figure 4D). According to previous studies, cigarette smoking affects oxidative protein folding [42] and the formation of the mitotic spindle in human organs [43], consistent with our findings.

Regarding results from the application of SR2 to the GBM data, BPs related to inflammation and immune response, neuron death, and myeloid cell differentiation are up-regulated in cells from recurrent GMB (Figure 4F). This is consistent with previous studies suggesting that aberrant activation of inflammatory responses is one main characteristic of GBM, and two other factors, cell death, and neuronal differentiation, in the GBM have been studied [44]. Moreover, recurrent GBM also shows higher levels of BPs related to inflammation and immune response compared with newly diagnosed GBM (Figure 4H).

### SR2 uncovers the effects of biological conditions on gene expression among different cell populations

While the previous section showed the overall effects of biological conditions over cell populations, it is more interesting to show the heterogenous effect of different biological conditions on gene expression for cell populations. Thus, phenotype- or disease-critical cell populations can be identified by comparing gene expression of cell populations across conditions, thus helping understand the mechanisms underlying phenotype (or disease).

Following the practice in one previous section in analyzing the expression of cell populations across biological conditions, adjusted expression profiles for cell populations under different biological conditions can be obtained by multiplying *W* with *V*. Then, we investigated the BPs enriched by genes in the upper or lower 5% quantile of the difference in the adjusted expression profile for the same cell population under different biological conditions. Then, the function analysis can uncover the effect of the biological condition on the cell population. For demonstration, we used the results from the applications of SR2 to the DM and COVID-19 dataset (shown in Figure 5).

**Figure 5.**
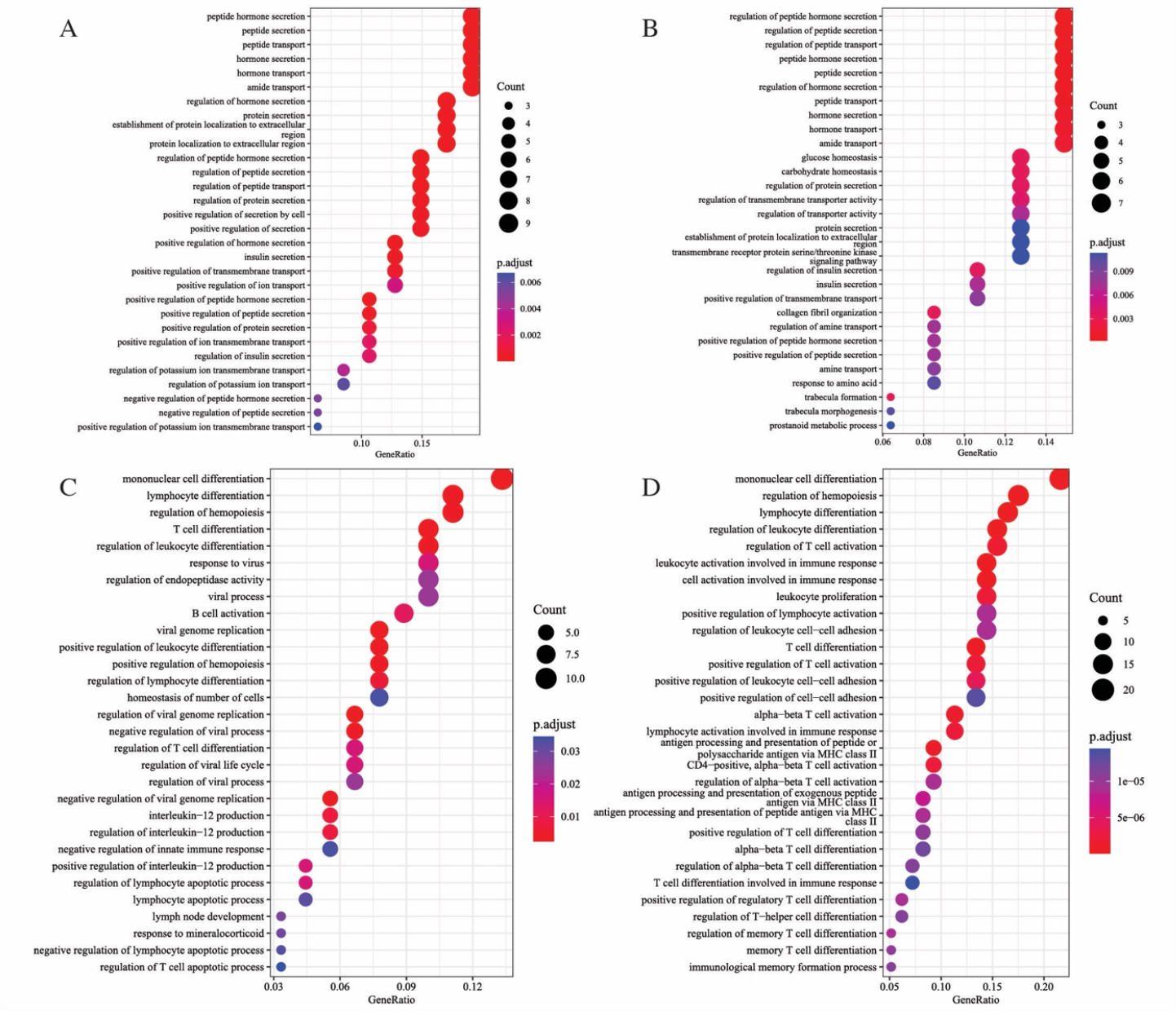
SR2 reveals the effects of biological conditions on gene expression for different cell types A. The figure shows the down-regulated BPs enriched by genes in the lower 5% quantile of the difference in the adjusted expression profiles for one of two larger beta cell populations between the diabetic and normal conditions. B. The figure shows the down-regulated BPs enriched by genes in the lower 5% quantile of the difference in the adjusted expression profiles for the other larger beta cell populations between the diabetic and normal conditions. C. The up-regulated BPs enriched by genes in the upper 5% quantile of the difference in the adjusted expression profiles for T cells between severe and light COVID-19 are displayed. D. The up-regulated BPs enriched by genes in the upper 5% quantile of the difference in the adjusted expression profiles for monocyte cells between severe and light COVID-19 are displayed.

Regarding the DM dataset, we conducted the above analysis to reveal the effects of diabetes on the gene expression for two major beta cell populations (Figures 5A and 5B), alpha cells (Figure S2C), PP cells (Figure S2D), stellate cells (Figure S2E). We observed that BPs revolved in insulin secretion are down-regulated in two beta populations from diabetic donors, compared with those from normal controls (Figure 5A and 5B). However, this finding is not observed for the other three cell types (Figures S2C, S2D, and S2E). This implies that beta cells are directly associated with diabetes. Moreover, the down-regulated BPs involved in peptide secretion (Figure S2E), glucose and carbohydrate homeostasis, amine transport, and peptide secretion (Figure S2C) are also enriched for stellate cells and alpha cells, from diabetic donors, respectively. This suggests that expression levels of genes involved in these BPs are lower in the two cell types from diabetic donors than from normal controls.

Subsequently, we revealed the effect of COVID-19 infection on gene expression for different cell populations with the COVID-19 dataset. Specifically, we uncovered the effects of severe COVID-19 infection on the gene expression of T cells (Figure 5C) and monocyte cells (Figure 5D). The BPs related to the differentiation of mononuclear cells, different kinds of T cells, and lymphocyte cells are up-regulated in monocyte cells from severe COVID-19 donors (Figure 5D). Similarly, levels of genes involved in these BPs are also high in T cells from severe COVID-19 donors (Figure 5C). Notably, the BPs involved in response to viral, such as response to viral and regulation of the viral life cycle and viral process, are also up-regulated in the T cells (Figure 5C). More importantly, levels of the BPs related to interleukin-2 production (Figure 5C), which plays a key role in the immune system, are also high in the T cells, not monocyte cells. This may suggest that monocyte cells promote the differentiation of T cells and that T cells play a key role in the immune response to severe COVID-19.

### SR2 reveals the molecular characteristics of donors

Here we demonstrate how SR2 can help reveal the molecular characteristics of donors in the application to the GBM dataset. First, we directly plotted the donor representations of 11 newly diagnosed GBM donors (Figure S1). Donors 01, 06, and 10 show obvious differences in molecular characteristics compared with the other eight donors (Figure S1). For demonstration purposes, we focus on discussing the molecular differences of these donors with metagenes 12, 13, and 15. Hence, we examined the BPs enriched by these metagenes. A joint analysis of Figure S1 and results from enrichment analysis suggest that donor 1 may suffer a poor progression of GBM, donor 6 may experience more severe hypoxia than other donors, and donor 10 has a stronger immune response and a lower level of gliogenesis and axonogenesis than other donors. Technical details of enrichment analysis and result interpretation are provided in SC.

However, due to a lack of clinical information on donors to validate our findings, further investigation is needed to explore our findings.

### Comparative study of SR2 on cell type annotation

Despite its interpretability demonstrated previously, SR2 also helps annotate cell types. To demonstrate its ability to annotate cell populations, we compared it against eight methods in cell identity annotation, including Seurat V3 [12], Liger [16], Harmony [14], BBKNN [8], ComBat [45], FastMNN [10], scINSIGHT [46], and Scanorama [13].

Practically, we followed the same procedure to ensure a fair comparison. After obtaining low-rank embeddings from each method, we performed cluster analysis with the low-rank embeddings using Louvain clustering [47] implemented in SCANPY [48]. We considered the number of cell types provided by the original data as the expected number of clusters. Thus, we adjusted the resolution until the expected number of clusters was attained. Then, we assessed the clustering performance for each method by Adjusted Rand Index (ARI), Adjusted Mutual Information (AMI), Normalized Mutual Information (NMI), and Homogeneity score. In the process, we considered the cell types annotated by original studies as true labels and ignored the unannotated observations. The cell types provided original studies are annotated by differential analysis with marker genes. Interested readers may refer to original studies for details.

To visualize cell representations from SR2, we employed UMAP [49] to obtain two-dimensional representations with low-rank embedding from SR2. In practice, we utilized the *umap* function with default settings from the R package *umap*. Then, we plotted the two-dimensional representations with points colored by annotated cell types.

### Comparison study of SR2 fitted on the whole data

First, we compared the performance of SR2 fitted on the whole data against those of the eight methods. In the comparison, we employed SR2 to three scRNA-seq datasets mentioned in the Datasets section. The result of the model comparison with the four metrics is presented in Figure 6A.

**Figure 6.**
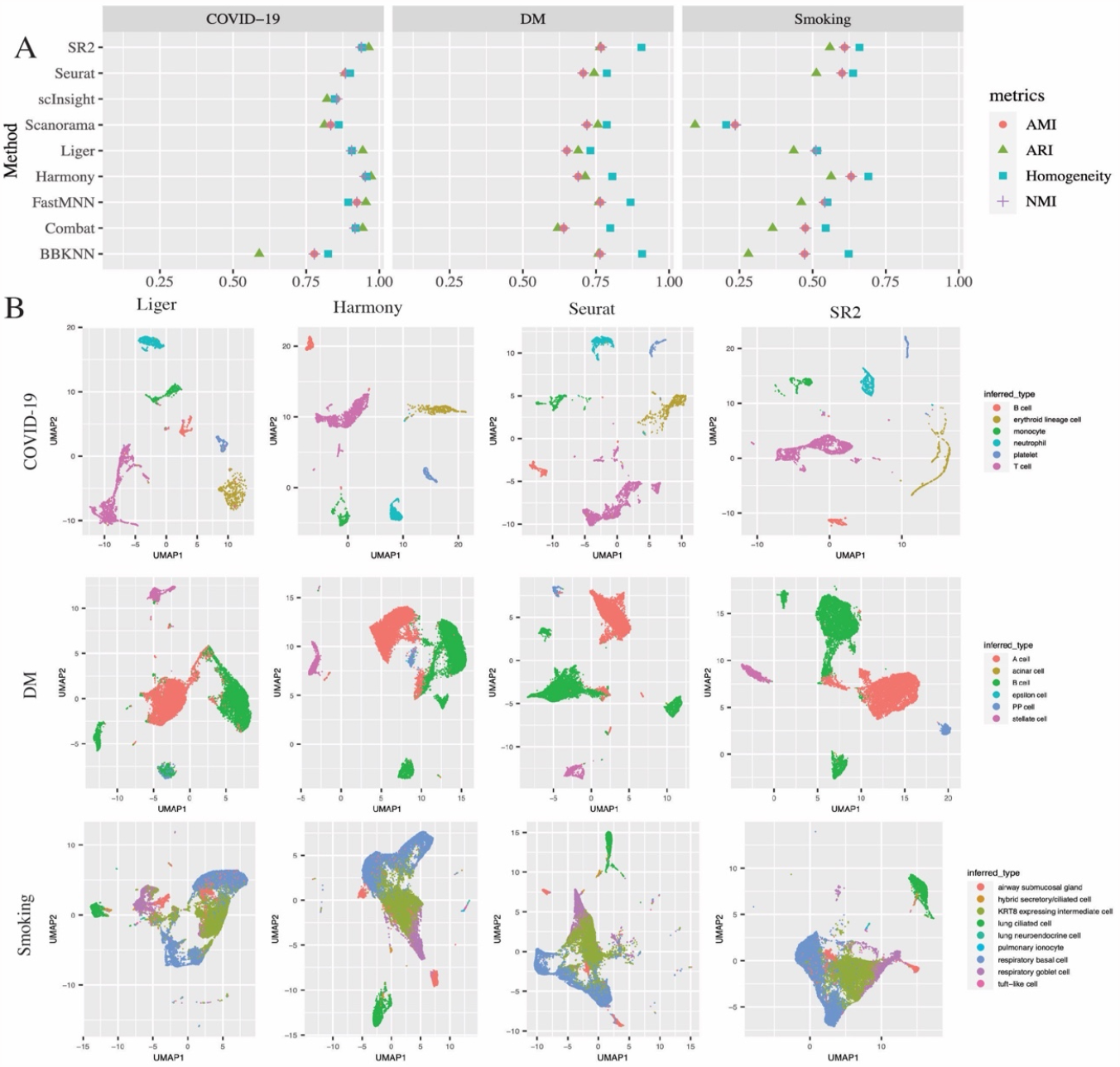
Comparative study of SR2 fitted with the whole data strategy A. The cluster performance of SR2 was compared to those of the other eight methods in terms of ARI, AMI, NMI, and Homogeneity score with the Smoking, DM, and COVID-19 datasets. The words “Smoking”, “DM”, and “COVID-19” stand for the Smoking, DM, and COVID-19 dataset, respectively. B. The figure visualizes two-dimensional representations from UMAP on low-rank embeddings from SR2 and the three recommended methods.

In general, across the four different evaluation metrics, SR2 achieved the start-of-the-art clustering performance (Figure 6A). Specifically, SR2 outperforms all other methods in the DM dataset in terms of ARI, NMI, and AMI (Supplementary ST1). In the other two datasets, SR2 achieved cluster performances comparable to the state-of-the-art methods (Figure 6A). In applications of scINSIGHT to the DM and Smoking datasets, it fails to yield results in two days, so we stop the program for time consideration.

To visually compare SR2 with the three methods (Liger, Seurat, and Harmony) recommended for scRNA-seq data integration by [50], we found that four methods demonstrate similar performance on the COVID-19 dataset (the first row of Figure 6B). In the analysis of the DM dataset, Harmony and SR2 outperform the other two methods, and SR2 seems to perform better in separating A (alpha) and B (beta) cells (the second row of Figure 6B). Specifically, Liger cannot well distinguish PP cells from B cells, and Seurat fails to distinguish PP cells from a subpopulation of A cells (the second row of Figure 1B). Moreover, quantitative measurement of these methods in separating A, B, and PP cells with silhouette scores also confirms our observation, with silhouette scores equal to 0.26, 0.22, 0.23, and 0.14 for SR2, Liger, Seurat, and Harmony, respectively. All methods perform similarly in the last application (the last row of Figure 6B), and this is also indicated by Figure 6A.

### Comparison study of batch SR2

We also compared the performance of batch SR2 on cell type annotation on the Immune and GBM datasets mentioned in the Datasets section.

The batch SR2 still achieves state-of-the-art performance in cell type annotations (Figure 7A). Specifically, batch SR2 outperforms all other methods in terms of ARI. Regarding the other three evaluation metrics, the performance of batch SR2 is similar to that of other methods. We also visually compared batch SR2 with the three recommended methods [50] in scRNA-seq data integration (shown in Figure 7B) and found that batch SR2 achieved a similar performance in cell type annotation in the comparison (Figure 7B).

**Figure 7.**
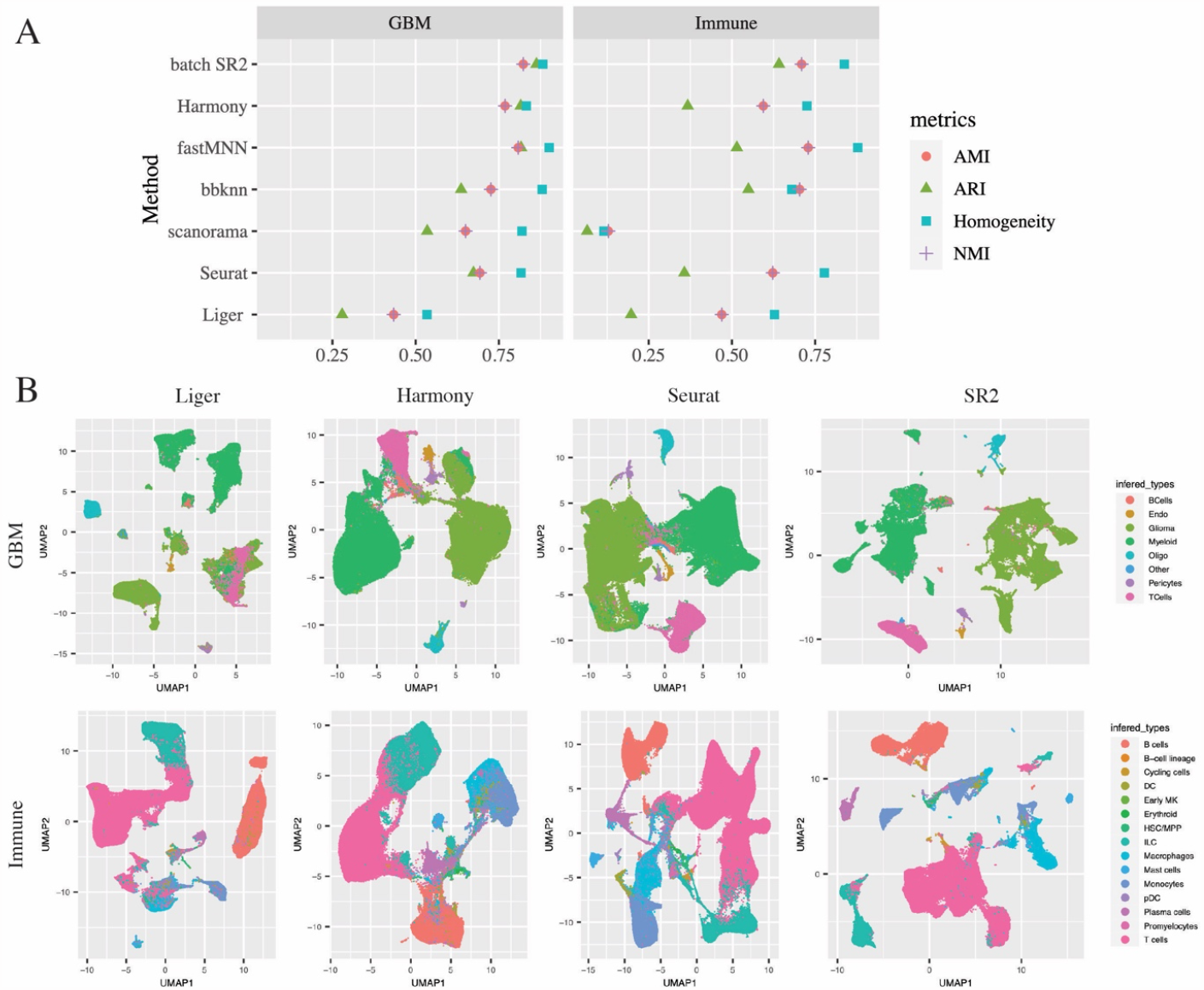
Comparative study of batch SR2 A. The cluster performance of SR2 was compared to those of the other eight methods in terms of ARI, AMI, NMI, and Homogeneity score on the GBM and Immune datasets. The performance of Combat and scINSIGHT are not shown in this Figure 7A and 7B due to its unsatisfactory scalability. B. Visualization of two-dimensional representations from UMAP on low-rank embeddings from batch SR2 and the three recommended methods.

In summary, SR2 achieves state-of-the-art performance in cell type annotation in our comparative studies, and the batch strategy improves the scalability of SR2 without impairing its performance in cell type annotation.

### Computational time and memory usage

SR2 is a computationally efficient framework for interpretable studies of scRNA-seq datasets. Though it is a complex computational tool, it is computationally efficient. The computational time for SR2 with the whole data strategy on the COVID-19 data, the DM dataset, and the Smoking dataset is ∼20 minutes, ∼1 hour, and ∼2.5 hours and its memory consumption in these analyses is less than 5G. For batch SR2, it takes roughly 6 hours and 12 G memory to complete the analyses with the GBM and Immune datasets, with the number of batches set to 20.

## Discussion

The broad applicability and potential of SR2 in scRNA-seq data analysis have been demonstrated through comprehensive applications. Specifically, SR2 with the whole data strategy was applied to three scRNA-seq datasets, and batch SR2 was studied in two scRNA-seq data of over 200 000 cells. The outputs from these applications enable various downstream analyses. Firstly, SR2 enables revealing changes in expression levels of genes across T2DM and control, COVID-19 outcomes, and GBM stages. The gene associated with these biological conditions were identified. Literature supports the relationship between the identified genes and the corresponding biological conditions. Secondly, it uncovers the BPs underlying T2DM, different COVID-19 outcomes, different smoking habits, and GBM stages, thus shedding light on disease mechanisms or the effect of biological conditions on gene expression among cells. Thirdly, it uncovers the effects of biological conditions on the expression of cell populations. For example, it reveals the effects of T2DM on gene expression for five major cell populations. Notably, the expression of genes involved in insulin secretion is down-regulated only in two beta cell populations from T2DM, compared to those from normal. This helps identify putative cell populations for diabetes. Fourthly, donor representations from SR2 characterize the molecular aspects of donors with GBM. Lastly, both SR2 fitted on whole data and batch SR2 achieve state-of-the-art performance in cell identity discovery.

In summary, SR2 has the following virtues, as demonstrated by our application studies: 1) SR2 is a general, flexible, and scalable framework for scRNA-seq data analysis, in which biological conditions can be defined upon study design and research question; 2) two different fitting strategies offered by SR2 make it tailored for analyzing scRNA-seq data of various sample size; 3) it models variation from biological conditions to enhance model interpretability and facilitate scRNA-seq data integration simultaneously; 4) it is extended to model the effect of biological conditions on gene expression for cell populations; 5) The output from SR2 enables various downstream analyses. Specifically, downstream analyses carried out include identifying phenotype- and disease-associated genes, revealing changes in gene expression across biological conditions, uncovering BPs underlying biological conditions (e.g., disease, treatment assignment), inferring phenotype-associated cell types, and revealing molecular characteristics of donors. A future direction of SR2 may include extending SR2 to analyze other single-cell omics.

To emphasize, SR2 focuses on model interpretability in scRNA-seq data analysis, though it achieves state-of-the-art performance in scRNA-seq data integration in terms of cell identity annotation. This interpretability aspect is crucial in computational methods for scRNA-seq data analysis, as many scRNA-seq studies aim to understand gene regulation across heterogeneous biological systems at the single-cell level [1], [3]-[5], [7], [51]. However, most existing methods for scRNA-seq data integration do not directly model variation from biological conditions. Thus, their ability to understand the effects of biological conditions at the single-cell level is limited. SR2 offers an solution to fill the gap in scRNA-seq data analysis. With a rapid rise in the scale of scRNA-seq data, SR2 further incorporates batch strategy to accommodate scRNA-seq data with arbitrary sample size. With these virtues and the emphasis, SR2 will benefit researchers in the biomedical field.

## Conclusion

This article proposed a computational framework, SR2, for single-cell RNA-seq data analysis. SR2 captures variation from heterogeneous biological conditions with matrix factorization and simultaneously decomposes the cellular variation with sparse representation learning. In SR2, we impose sparse regularization on embeddings of cells and restrict the norm of each component in the sparse representation learning. SR2 is a general and flexible framework in which biological conditions can be defined based on the study aim. The outputs from SR2 provides informative inputs for various downstream analysis, such as revealing gene activities across the biological condition, identifying phenotype-associated genes, and uncovering BPs underlying biological conditions. Moreover, SR2 can also uncover the effects of biological conditions on gene expression for cell populations, thus identifying putative cell populations associated with biological conditions. Its wide applicability in biomedical data analysis has been demonstrated through comprehensive applications. Further extensions include analysis of other single-cell omics.

## Methods

Here we propose a novel statistical approach, SR2, to decompose cellular variations of scRNA-Seq samples with considering variation from multiple biological conditions (e.g., donor, tissue, phenotype) into low-rank latent spaces to facilitate downstream analysis. In SR2, we integrate matrix factorization, which is utilized to capture variation from biological conditions, and sparse representation learning to obtain embeddings of cells. In employing sparse representation learning, we introduce sparse regularization on embeddings of cells to facilitate cell population discovery and norm constraint on each component of gene representations to ensure equal scale.

### The SR2 model and its extension

For illustration, let *Z*^*N*×*M*^ denote the matrix of log-normalized scRNA-Seq expression levels of *N* samples of *M* genes. The *N* samples can originate from several biological conditions (e.g., individuals, phenotypes, tissues, or disease phases). Here we use donor and phenotype as an example for demonstration. The samples come from *N*_1_ donors with *N*_2_ phenotypes. Thus, the expression level of gene *m*for sample *i* obtained from donor *j* with phenotype *t, z*_*im*_, can be modeled as

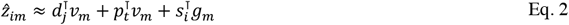

where *d*_*j*_, *p*_*t*_, *ν*_*m*_ are vectors of length *K*_1_, *s*_*i*_, *g*_*m*_ are vectors of length *K*_2_. The donor and phenotype information of the samples is known. The rationale behind the modeling is that different donors demonstrate heterogeneity in gene expression via pathways and that different phenotypes (or conditions) affect donors’ gene expression via the same pathways. In practice, we also found that increasing model complexity by modeling variation from donor and phenotype in two separate latent spaces does not lead to explain more variation from them. Therefore, we capture variation from donors and phenotypes by restricting in a shared low-rank latent space. An extra benefit of this practice is that it can reduce model complexity and improve optimization efficiency without impairing model interpretability. Intuitively, after modeling variation from biosamples (e.g., donor and phenotypes), we capture variation across different scRNA-Seq samples in another low-rank latent space. This practice model the cellular variation in a more independent manner and increase model flexibility, compared with restricting our modeling in only one latent space. In our implementation, we considered the option to restrict *ν*_*m*_ and *g*_*m*_ to be the same. Thus, the last term on the right of Eq. 2 helps obtain low-rank embeddings for different cells. Moreover, in SR2, biological conditions can be defined based on the research questions and study design.

The objective function for Eq. 2 is formulated as

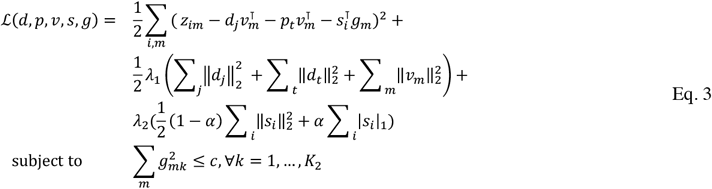

In the above equation, *g*_*mk*_ is the *k*-th element of *g*_*m*_, and *c* is a constant (usually 1). We introduce the elastic net penalty on the cell representation *s*_*i*_ to encourage sparsity to facilitate cell clustering. Moreover, the norm constraint is imposed to ensure the same scale for each component in decomposing cellular variation.

Equation 3 can be represented with matrix operation. Let 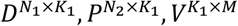 are matrices of latent representations of *N*_1_ donors, *N*_2_ phenotypes, and *M* genes, respectively, and denote 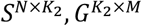 latent representations for *N* cells and *M* genes after modeling variation from donor and phenotype. The matrix representation of Equation 2 can be written as follows:

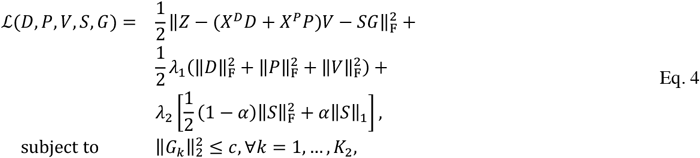

where *X*^*D*^, *X*^*P*^ are indicator matrices of *N* rows and *N*_1_ and *N*_2_ colunns, respectively, which are the dummy variables for *N* samples, and *G*_*k*_ is the *k*-th row of matrix *G. c* is a constant, restricting the scale of each component of *G*.

The above formulation of SR2 can be extended to explore the effects of biological conditions on gene expression for cell populations after cell type annotation. To exploit this idea, we model the expression level of gene *m* for sample *i* from cell population *k* and obtained from donor *j* with phenotype *t, z*_*im*_ as

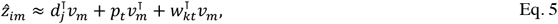

where *d*_*j*_, *p*_*t*_, *w*_*kt*_, *ν*_*m*_ are vectors of length *K*, and *w*_*kt*_ is the latent representation for cell population *k* from donors with phenotype *t*, which captures the interactive effect between phenotypes and cell populations while controlling variation from donors and phenotypes. In the above equation, the biological condition and cell populations can be defined based on research questions, and other biological conditions (e.g., tissue or treatment assignment) can also be incorporated. Thus, the extension provides a useful tool for exploring the effects of biological conditions on the expression behavior of cell populations obtained from other computational tools. Throughout this study, we name the new formulation of SR2 as extended SR2.

The objective function for the above equation with matrix notation is defined as

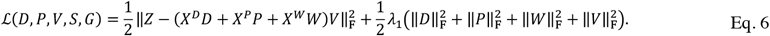

Here *X*^*W*^ and 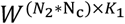 denote the indicator matrix of *N* rows and *N*_*3*_ columns and latent representations for *N*_c_ cell populations under *N*_2_ biological conditions, respectively. Other notations are the same as in Eq. 4.

### Model fitting

Alternating block coordinate descent (BCD) was employed to optimize Equation 2. Practically, each time we update a set of independent parameters in parallel with all other parameters fixed. In each iteration of BCD, we update each set of parameters of our model sequentially and repeat the process until the stopping criteria meets.

#### Optimize with the whole data strategy

With the objective function and notations defined in Eq. 4 and 6, we can easily derive closed forms for updating *d*_*j*_, *p*_*t*_, *ν*_*m*_. Specifically, we have the following update for *ν*_*m*_

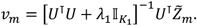

For the objective function defined by Eq. 4, 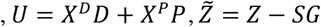, and 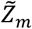 is the *m*-th column of 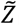. However, *U* = *X*^*D*^*D* + *X*^*P*^*P* + *X*^*W*^*W*, and 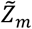 is the *m*-th column of 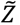, where 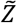 is equal to *Z*, when the objective function is defined by Eq. 6. Similarly, the update for *d*_*j*_ is

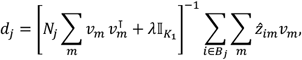

where 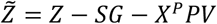 and 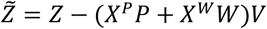 for the objective function defined by Eq. 4 and Eq. 6, respectively, *B*_*j*_ is the set of indices of samples from donor *j*, and *N*_*j*_ is the number of elements in *B*_*j*_. Likewise, the update for *p*_*t*_ is

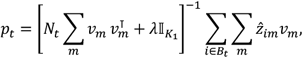

where 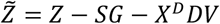 and 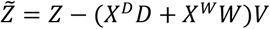 for the objective function defined by Eq. 4 and Eq. 6, respectively, *B*_*t*_ is the set of indices of samples from donors with phenotype *t*, and *N*_*t*_ is the number of elements in *B*_*t*_. Finally, the update for *w*_*kt*_ is

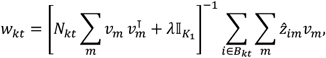

where 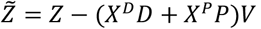 for the objective function defined by Eq. 6, respectively, *B*_.*t*_ is the set of indices of samples of cell type *k* from donors with phenotype *t*, and *N*_*kt*_ is the number of elements in *B*_*t*_.

When optimizing Eq. 4 with respect to *G*, the Lagrange dual proposed in the study [52] is employed. The Lagrange dual for our problem is of the following form

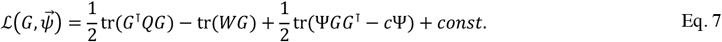

Here 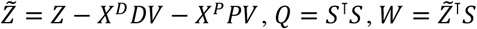, and Ψ is a *K*_2_ × *K*_2_ diagonal matrix with dual variables *ψ* expanding along its diagonal. By taking the derivative with respect to *G*, we have

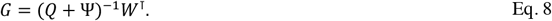

Then, by substituting Eq. 8 into the Lagrange dual above, we have the following dual for our problem

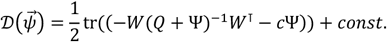

The gradient ∇ and Hessian *H* for 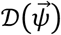 with respect to 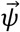 can be derived as follows:

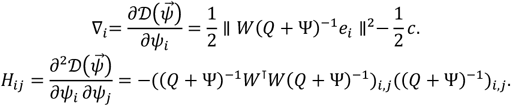

Newton’s method is used to optimize our dual problem with respect to Ψ. Thus, the update for Ψ at iteration *t* can be written as

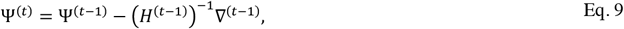

where Ψ^(*t*−1)^, *H*^(*t*−1)^, ∇^(*t*−1)^ are the diagonal matrix of *ψ*, gradient, and Hessian matrix at iteration *t* − 1. In practice, we alternatively compute the updates for *G*, Ψ until the sum of the squared difference in Ψ between two consecutive iterations less than a predefined threshold (10^−4^ is used in our studies). Further details on the derivation of Lagrange dual are provided in the Supplementary.

When optimizing Equation 4 with respect to *s*_*i*_, the *i*-th row of *S*, our objective function can be simplified to

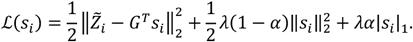

Here 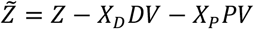, and 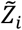 is the vector of the *i*-th row of 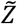. To solve the problem, random coordinate descent (RCD) with strong rules is employed, which was proposed in our previous study [53].

#### Optimize with the batch strategy

When the sample size of data is huge, optimizing SR2 with the whole data is memory-demanding and computationally intensive. To relieve this issue, we propose a batch strategy to optimize SR2. Practically, we split the whole data into several batches and optimized our object function with one batch each time to lower the memory consumption and computation burden.

The key to performing batch optimization of SR2 is to come up with a *surrogate* that asymptotically converges to the same solution defined by Equation 4. As inspired by one previous study [54], we define the following *surrogate* for our objective

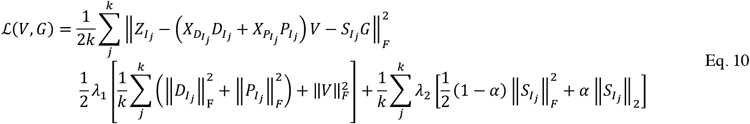

Here *k* is the number of batches, *I*_*j*_ denote the set of indices of cells from batch *j*, and 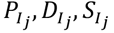 denote the representation for phenotypes, donors, and cells that are obtained from batch *j*. Technically, the *surrogate* for Eq. 6 is a subproblem defined by Eq. 10, so we will focus on the technical details of our objective defined by Eq. 10 and not diverge to discuss the details of the *surrogate* for Eq. 6, which can be easily solved by Algorithm 1 with subtle changes. Algorithm 1 below is proposed to optimize the *surrogate* defined by Eq. 10.

##### Algorithm 1: Batch SR2

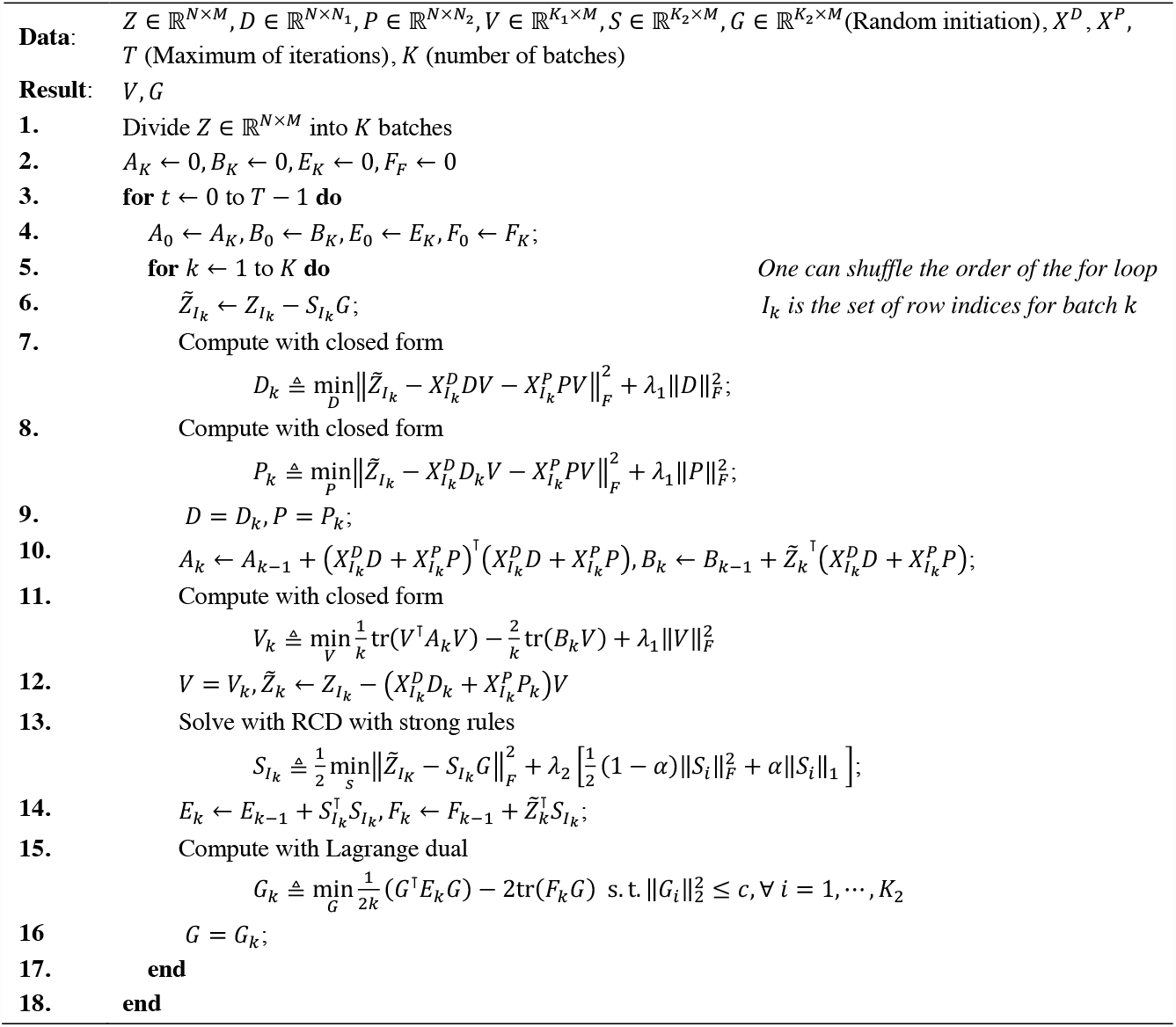

In Algorithm 1, we notice that *A*_*k*_, *B*_*k*_, *E*_*k*_, *F*_*k*_ carry all information for the same batch from all previous iterations. Actually, the information from early iterations is outdated. Mairal et al. [54] suggested accelerating convergence by removing old information for the same batch from these matrices. Specifically, owning to the design of our algorithm, we use the following equations to exploit this idea:

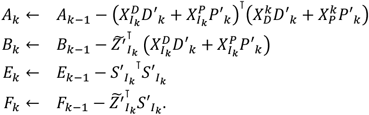

Here *D*′_*k*_, *P*′_.*k*._, *S*′_*k*_ are the matrices for donor, phenotype, and cell representations for batch *k* from iteration *t* − 1. With slight abuse of notation, 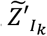 in the second and fourth line of the above equation are computed with equations in lines 12 and 14 in Algorithm 1, respectively.

### Initialization, hyperparameter tuning, and the stopping criteria

In SR2, all latent variables (*D, P, V, S, G*) are initiated from the normal distribution *N*(0,0.001). For initiating *s*_*i*_ in by Eq. 10 in optimization, we use the solution for *s*_*i*_ from the previous iteration.

For model selection in SR2, grid search is utilized to select hyperparameters *λ*_1_, *λ*_2_, *α, K*_1_, *K*_2_. In practice, when the number of observations is huge (e.g., ≥ 500,000), we randomly and evenly draw a small proportion (e.g., 0.1) of scRNA-Seq samples as a dataset for model selection. Then, we randomly draw 1c*t* of elements from the dataset matrix as a test set. For each combination of candidate hyperparameters, we run alternating BCD several iterations (e.g., 20) and choose the one with the best performance on the test set in terms of root-mean-square error (RMSE) on the test set.

In practice, we found that SR2 favors the Lasso penalty in applications. Thus, we set *α* equal to 1 to simplify the model selection. Meanwhile, our two hyperparameters *K*_1_ and *λ*_1_ are somewhat redundant since one can increase *K*_1_ and *λ*_1_ simultaneously without changing model complexity. Therefore, we set *K*_1_ = *K*_2_ in the model selection.

The detailed procedure for model selection is as follows. First, we set *λ*_1_, *λ*_2_ to a small number (e.g., c.1) to avoid singularity in matrix inverse, and *α* is fixed to 1, and choose the ranks of latent representations *K*_1_ = *K*_2_ from sequences from 10 to 40 with step size 2 with grid search. After choosing the ranks of latent dimensions, we define a broad parameter grid for *λ*_1_, *λ*_2_ and perform a grid search. One may also refine the parameter grid based on the performance of our parameter sets on the test set. Finally, we select the parameters *λ*_1_, *λ*_2_ with the best performance on the test set and run SR2 with the selected parameters until the stopping criteria meet.

In our study, the stopping criteria are defined as

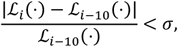

where ℒ_*i*_(⋅) is the loss in the *i*-th iteration and calculated every ten iterations to reduce the computational burden, and *σ* is a predefined threshold and set to 10^−8^ in our experiment.

## Supporting information

Supplementary Content

## Acknowledgement

This work has been supported by the Chinese University of Hong Kong startup grant (4930181), the Chinese University of Hong Kong Science Faculty’s Direct Grant for Research 2022/2023, the Chinese University of Hong Kong Science Faculty’s Collaborative Research Impact Matching Scheme (CRIMS 4620033) and Hong Kong Research Grant Council (GRF 14301120, GRF 14300923).

## Declarations

The authors declare that they have no competing interests.

## Notes

https://github.com/kai0511/SR2

### Competing Interest Statement

The authors have declared no competing interest.

## References

[1] C. Domínguez Conde et al, “Cross-tissue immune cell analysis reveals tissue-specific features in humans,” Science, vol. 376, (6594), pp. eabl5197, 2022.

[2] Z. Fang et al, “Single-cell heterogeneity analysis and CRISPR screen identify key β-cell-specific disease genes,” Cell Reports, vol. 26, (11), pp. 3132–3144. e7, 2019.

[3] A. Silvin et al, “Elevated calprotectin and abnormal myeloid cell subsets discriminate severe from mild COVID-19,” Cell, vol. 182, (6), pp. 1401–1418. e18, 2020.

[4] N. Abdelfattah et al, “Single-cell analysis of human glioma and immune cells identifies S100A4 as an immunotherapy target,” Nature Communications, vol. 13, (1), pp. 767, 2022.

[5] K. C. Goldfarbmuren et al, “Dissecting the cellular specificity of smoking effects and reconstructing lineages in the human airway epithelium,” Nature Communications, vol. 11, (1), pp. 1–21, 2020.

[6] J. D. Welch et al, “Single-cell multi-omic integration compares and contrasts features of brain cell identity,” Cell, vol. 177, (7), pp. 1873–1887. e17, 2019.

[7] L. Jerby-Arnon et al, “A cancer cell program promotes T cell exclusion and resistance to checkpoint blockade,” Cell, vol. 175, (4), pp. 984–997. e24, 2018.

[8] K. Polański et al, “BBKNN: fast batch alignment of single cell transcriptomes,” Bioinformatics, vol. 36, (3), pp. 964–965, 2020.

[9] J. Ming et al, “FIRM: fast Integration of singlecell RNA-sequencing data across multiple platforms,” bioRxiv, 2020.

[10] L. Haghverdi et al, “Batch effects in single-cell RNA-sequencing data are corrected by matching mutual nearest neighbors,” Nat. Biotechnol., vol. 36, (5), pp. 421–427, 2018.

[11] K. Qian et al, “scINSIGHT for interpreting single-cell gene expression from biologically heterogeneous data,” Genome Biol., vol. 23, (1), pp. 1–23, 2022.

[12] A. Butler et al, “Integrating single-cell transcriptomic data across different conditions, technologies, and species,” Nat. Biotechnol., vol. 36, (5), pp. 411–420, 2018.

[13] B. Hie, B. Bryson and B. Berger, “Efficient integration of heterogeneous single-cell transcriptomes using Scanorama,” Nat. Biotechnol., vol. 37, (6), pp. 685–691, 2019.

[14] I. Korsunsky et al, “Fast, sensitive and accurate integration of single-cell data with Harmony,” Nature Methods, vol. 16, (12), pp. 1289–1296, 2019.

[15] T. Stuart et al, “Comprehensive integration of single-cell data,” Cell, vol. 177, (7), pp. 1888–1902. e21, 2019.

[16] J. D. Welch et al, “Single-cell multi-omic integration compares and contrasts features of brain cell identity,” Cell, vol. 177, (7), pp. 1873–1887. e17, 2019.

[17] A. Butler et al, “Integrating single-cell transcriptomic data across different conditions, technologies, and species,” Nat. Biotechnol., vol. 36, (5), pp. 411–420, 2018.

[18] B. Hie, B. Bryson and B. Berger, “Efficient integration of heterogeneous single-cell transcriptomes using Scanorama,” Nat. Biotechnol., vol. 37, (6), pp. 685–691, 2019.

[19] G. L. Stein-O’Brien et al, “Enter the matrix: factorization uncovers knowledge from omics,” Trends in Genetics, vol. 34, (10), pp. 790–805, 2018.

[20] K. A. Jagadeesh et al, “Identifying disease-critical cell types and cellular processes by integrating single-cell RNA-sequencing and human genetics,” Nat. Genet., vol. 54, (10), pp. 1479–1492, 2022.

[21] C. Boutari et al, “Humanin and diabetes mellitus: A review of in vitro and in vivo studies,” World Journal of Diabetes, vol. 13, (3), pp. 213, 2022.

[22] I. Garin et al, “Recessive mutations in the INS gene result in neonatal diabetes through reduced insulin biosynthesis,” Proceedings of the National Academy of Sciences, vol. 107, (7), pp. 3105–3110, 2010.

[23] P. Kaur et al, “SERPINE 1 links obesity and diabetes: a pilot study,” Journal of Proteomics & Bioinformatics, vol. 3, (6), pp. 191, 2010.

[24] S. I. Alsters et al, “Truncating homozygous mutation of carboxypeptidase E (CPE) in a morbidly obese female with type 2 diabetes mellitus, intellectual disability and hypogonadotrophic hypogonadism,” PloS One, vol. 10, (6), pp. e0131417, 2015.

[25] V. Alur et al, “Analysis of key genes and pathways associated with the pathogenesis of Type 2 diabetes mellitus,” bioRxiv, pp. 2021.08. 12.456106, 2021.

[26] S. Costes et al, “UCHL1 deficiency exacerbates human islet amyloid polypeptide toxicity in β-cells: evidence of interplay between the ubiquitin/proteasome system and autophagy,” Autophagy, vol. 10, (6), pp. 1004–1014, 2014.

[27] E. Kvedaraite et al, “Major alterations in the mononuclear phagocyte landscape associated with COVID-19 severity,” Proceedings of the National Academy of Sciences, vol. 118, (6), pp. e2018587118, 2021.

[28] J. S. Muhammad et al, “SARS-CoV-2-induced hypomethylation of the ferritin heavy chain (FTH1) gene underlies serum hyperferritinemia in severe COVID-19 patients,” Biochem. Biophys. Res. Commun., vol. 631, pp. 138–145, 2022.

[29] T. Wu et al, “Expression of ferritin light chain (FTL) is elevated in glioblastoma, and FTL silencing inhibits glioblastoma cell proliferation via the GADD45/JNK pathway,” PloS One, vol. 11, (2), pp. e0149361, 2016.

[30] V. Ravi et al, “Liposomal delivery of ferritin heavy chain 1 (FTH1) siRNA in patient xenograft derived glioblastoma initiating cells suggests different sensitivities to radiation and distinct survival mechanisms,” PLoS One, vol. 14, (9), pp. e0221952, 2019.

[31] H. M. Said et al, “GAPDH is not regulated in human glioblastoma under hypoxic conditions,” BMC Molecular Biology, vol. 8, (1), pp. 1–13, 2007.

[32] A. Zottel et al, “Anti-Vimentin Nanobody Decreases Glioblastoma Cell Invasion In Vitro and In Vivo,” Cancers, vol. 15, (3), pp. 573, 2023.

[33] R. Huang et al, “Retinol binding protein 4 impairs pancreatic beta-cell function, leading to the development of type 2 diabetes,” Diabetes, vol. 67, (Supplement_1), 2018.

[34] D. Flamez et al, “A genomic-based approach identifies FXYD domain containing ion transport regulator 2 (FXYD2) γa as a pancreatic beta cell-specific biomarker,” Diabetologia, vol. 53, pp. 1372–1383, 2010.

[35] A. Mahajan et al, “Fine-mapping type 2 diabetes loci to single-variant resolution using high-density imputation and islet-specific epigenome maps,” Nat. Genet., vol. 50, (11), pp. 1505–1513, 2018.

[36] S. C. Bearrows et al, “Chromogranin B regulates early-stage insulin granule trafficking from the Golgi in pancreatic islet β-cells,” J. Cell. Sci., vol. 132, (13), pp. jcs231373, 2019.

[37] M. Tewari et al, “Lymphoid expression and regulation of A20, an inhibitor of programmed cell death.” Journal of Immunology (Baltimore, Md.: 1950), vol. 154, (4), pp. 1699–1706, 1995.

[38] D. Ramljak et al, “Early response of CD8 T cells in COVID-19 patients,” Journal of Personalized Medicine, vol. 11, (12), pp. 1291, 2021.

[39] H. C. Greenwood, S. R. Bloom and K. G. Murphy, “Peptides and their potential role in the treatment of diabetes and obesity,” The Review of Diabetic Studies, vol. 8, (3), 2011.

[40] S. J. Brandt et al, “Gut hormone polyagonists for the treatment of type 2 diabetes,” Peptides, vol. 100, pp. 190–201, 2018.

[41] S. André et al, “T cell apoptosis characterizes severe Covid-19 disease,” Cell Death & Differentiation, pp. 1–14, 2022.

[42] H. Kenche et al, “Cigarette smoking affects oxidative protein folding in endoplasmic reticulum by modifying protein disulfide isomerase,” The FASEB Journal, vol. 27, (3), pp. 965–977, 2013.

[43] M. T. Landi et al, “Gene expression signature of cigarette smoking and its role in lung adenocarcinoma development and survival,” PloS One, vol. 3, (2), pp. e1651, 2008.

[44] P. Guichet et al, “Cell death and neuronal differentiation of glioblastoma stem-like cells induced by neurogenic transcription factors,” Glia, vol. 61, (2), pp. 225–239, 2013.

[45] W. E. Johnson, C. Li and A. Rabinovic, “Adjusting batch effects in microarray expression data using empirical Bayes methods,” Biostatistics, vol. 8, (1), pp. 118–127, 2007.

[46] K. Qian et al, “scINSIGHT for interpreting single-cell gene expression from biologically heterogeneous data,” Genome Biol., vol. 23, (1), pp. 1–23, 2022.

[47] V. D. Blondel et al, “Fast unfolding of communities in large networks,” Journal of Statistical Mechanics: Theory and Experiment, vol. 2008, (10), pp. P10008, 2008.

[48] F. A. Wolf, P. Angerer and F. J. Theis, “SCANPY: large-scale single-cell gene expression data analysis,” Genome Biol., vol. 19, (1), pp. 1–5, 2018.

[49] L. McInnes, J. Healy and J. Melville, “Umap: Uniform manifold approximation and projection for dimension reduction,” arXiv Preprint arXiv:1802.03426, 2018.

[50] H. T. N. Tran et al, “A benchmark of batch-effect correction methods for single-cell RNA sequencing data,” Genome Biol., vol. 21, (1), pp. 1–32, 2020.

[51] M. B. Johnson et al, “Single-cell analysis reveals transcriptional heterogeneity of neural progenitors in human cortex,” Nat. Neurosci., vol. 18, (5), pp. 637–646, 2015.

[52] H. Lee et al, “Efficient sparse coding algorithms,” Advances in Neural Information Processing Systems, vol. 19, 2006.

[53] Z. Kai et al, “INSIDER: Interpretable Sparse Matrix Decomposition for Bulk RNA Expression Data Analysis,” bioRxiv, 2022.

[54] J. Mairal et al, “Online dictionary learning for sparse coding,” in Proceedings of the 26th Annual International Conference on Machine Learning, 2009,.

