## Supplementary Content for "SR2: Sparse Representation Learning for Scalable Single-cell RNA Sequencing Data Analysis"

### Lagrange dual for learning bases

Here we derive the updates of $G$ in the problem defined in the following:

$$\arg\min_{G} \frac{1}{2}\left\| Z-SG \right\|_{F}^{2}+\frac{1}{2}\left( 1-\alpha\right)\lambda\left\| S \right\|_{F}^{2}+\lambda\alpha\left| S \right|_{1} subject to \parallel G_{i}\parallel_{2}^{2}\leq1,\forall i=1,\ldots,k,$$

where $Z$ is the matrix to be approximated, $S$ is the sparse representation, and $G$ is the matrix for the dictionary we seek to obtain. In the equation, $G_{i}$ is the $i$-th row of $G$. To solve the problem, we consider the following Lagrange:

$$\mathcal{L}\left( G,\vec{\lambda} \right)=\frac{1}{2}\mathrm{tr}\left( \left( Z-SG \right)^{⊺}\left( Z-SG \right) \right)+\frac{1}{2}\mathrm{tr}\left( \Lambda GG^{⊺}-\Lambda\right),$$

where each $\lambda_{k}\geq0$ is a dual variable and $\Lambda=diag\left( \vec{\lambda} \right)$. Taking derivative with respect to $G$, we obtain

$$\frac{\partial\mathcal{L}\left( G,\vec{\lambda} \right)}{\partial G}=-Z^{⊺}S+S^{⊺}SG+\Lambda G=0.$$

Here $F=Z^{⊺}S$ and $E=S^{⊺}S$. With these notations, we have

$$G=(E+\Lambda)^{-1}F^{⊺}.$$

Eq. 1

Substituting Equation 9 into the Lagrange defined above, we can further derive the Lagrange dual for our problem:

$$\begin{matrix} \mathcal{D}\left( \vec{\lambda} \right)= & \min_{G}\mathcal{L}\left( G,\vec{\lambda} \right)=\frac{1}{2}tr(\left( Z^{⊺}Z-2FG+G^{⊺}EG+\Lambda GG^{⊺}-\Lambda\right)) \\ = & \frac{1}{2}tr((Z^{⊺}Z-2F(E+\Lambda)^{-1}F^{⊺}+F(E+\Lambda)^{-1}E(E+\Lambda)^{-1}F^{⊺}+ \\ & F(E+\Lambda)^{-1}\Lambda(E+\Lambda)^{-1}F^{⊺}-\Lambda)) \\ = & \frac{1}{2}tr(\left( Z^{⊺}Z-2F(E+\Lambda)^{-1}F^{⊺}+F(E+\Lambda)^{-1}F^{⊺}-\Lambda\right)) \\ = & \frac{1}{2}tr(\left( Z^{⊺}Z-F(E+\Lambda)^{-1}F^{⊺}-\Lambda\right)), \end{matrix}$$

which can be further formulated as

$$\mathcal{D}\left( \vec{\lambda} \right)=\frac{1}{2}tr(\left( Z^{⊺}Z-F(E+\Lambda)^{-1}F^{⊺}-\Lambda\right)).$$

The gradient and Hessian of $\mathcal{D}\left( \vec{\lambda} \right)$ are computed as follows:

$$\frac{\partial\mathcal{D}\left( \vec{\lambda} \right)}{\partial\lambda_{i}}=\frac{1}{2}\left\| F\left( E+\Lambda\right)^{-1}e_{i} \right\|^{2}-\frac{1}{2},$$

$$\frac{\partial^{2}\mathcal{D}\left( \vec{\lambda} \right)}{\partial\lambda_{i}\partial\lambda_{j}}=-(\left( E+\Lambda\right)^{-1}F^{⊺}F\left( E+\Lambda\right)^{-1})_{i,j}(\left( E+\Lambda\right)^{-1})_{i,j}.$$

The Newton’s method is used to optimize the above problem.

### SR2 reveals the molecular characteristics of donors

Here we demonstrate how SR2 can help reveal the molecular characteristics of donors in the application to the GBM dataset. First, we directly plotted the donor representations of 11 newly diagnosed GBM donors (shown in Figure S1). We see that the three donors (01, 06, and 10) show obvious differences in molecular characteristics, compared with the other eight donors (Figure S1). For demonstration purposes, we focus on discussing the molecular differences of these donors with metagenes 12, 13, and 15 (highlighted in Figure S1), though some other metagenes may help reveal the difference.

The loading of the 13^th^ and 15^th^ metagenes is highly positive for donor 10. Subsequent enrichment analysis on the expression profile of metagenes 13 and 15 reveals that they encode genes up-regulating inflammation responses (Figure S3C) and T cell activation (Figure S3D), respectively, and down-regulating gliogenesis and glial cell differentiation (Figure S3E and S3F). However, the loading of the 15^th^ metagene for donor 6 is negative, suggesting the level of inflammation response of the donor is low. Moreover, metagene 12 encodes genes up-regulating response to hypoxia (Figure S3A) and down-regulating peptidase and endopeptidase activities (Figure S3B), which is reported to promote glioma malignancy [40] and tumorigenesis [41]. Thus, our finding suggests that donor 1 may suffer a poor progression of GBM, donor 6 may experience more severe hypoxia than other donors, and donor 10 has a strong immune response and a lower level of gliogenesis and axonogenesis. Note that for demonstration purposes we only discussed the molecular characteristics of three donors via enrichment analysis of three metagenes, so a more comprehensive analysis is needed to reveal the big picture of the molecular characteristics of donors.

Figure S1 heatmap of donor representation for 11 newly diagnosed GBM

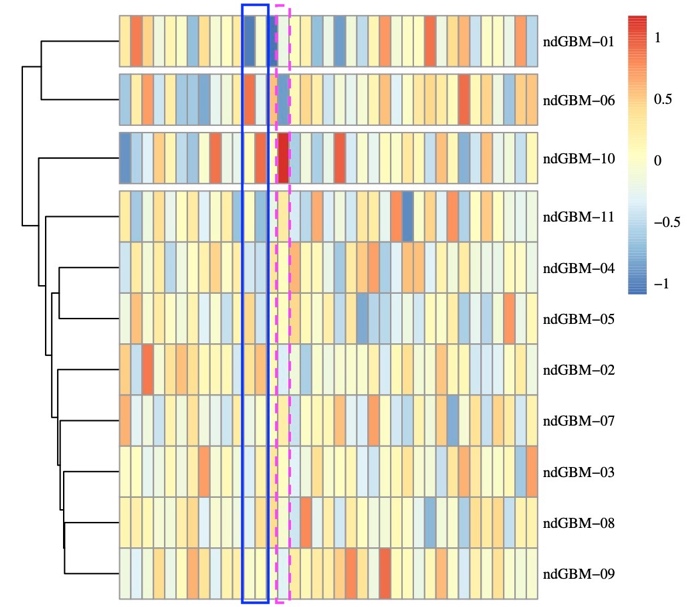

Figure S2 supplementary figures from analysis of results from SR2 in application to the DM dataset

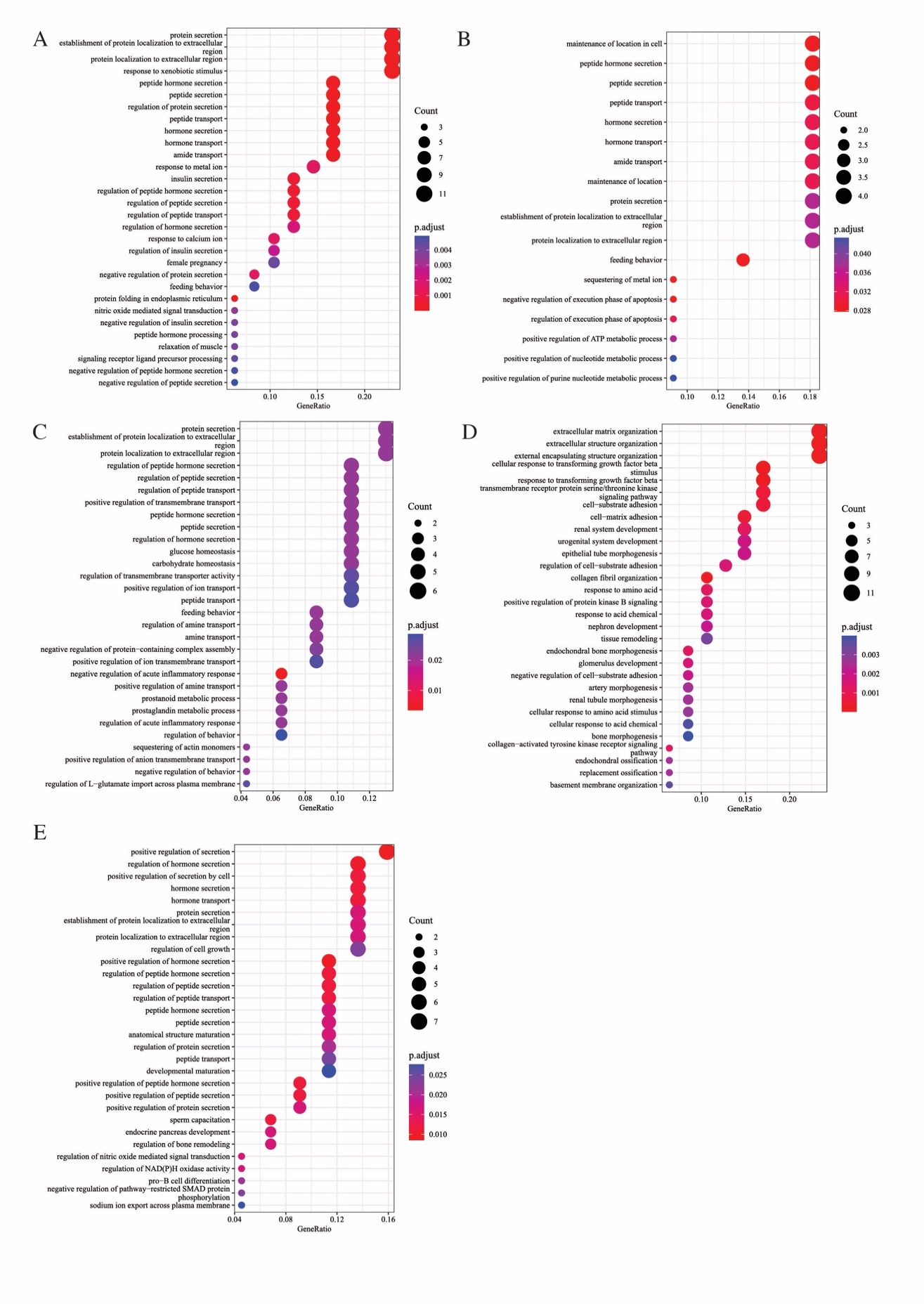

A. The figure displays the top up-regulated 30 BPs enriched by top genes in the adjusted expression profiles for another large beta cell population from normal donors in the analysis of the DM dataset.
B. The figure displays the top up-regulated 30 BPs enriched by top genes in the adjusted expression profiles for the smaller beta cell population from normal donors in the analysis of the DM dataset.
C. The down-regulated BPs enriched by genes in the upper 5% quantile of the difference in the adjusted expression profiles for alpha cells between the diabetic and normal conditions are displayed.
D. The down-regulated BPs enriched by genes in the upper 5% quantile of the difference in the adjusted expression profiles for PP cells between the diabetic and normal conditions are displayed.
E. The down-regulated BPs enriched by genes in the upper 5% quantile of the difference in the adjusted expression profiles for stellate cells between the diabetic and normal conditions are displayed.

Figure S3 supplementary figures from analysis of results from SR2 in application to the GBM dataset

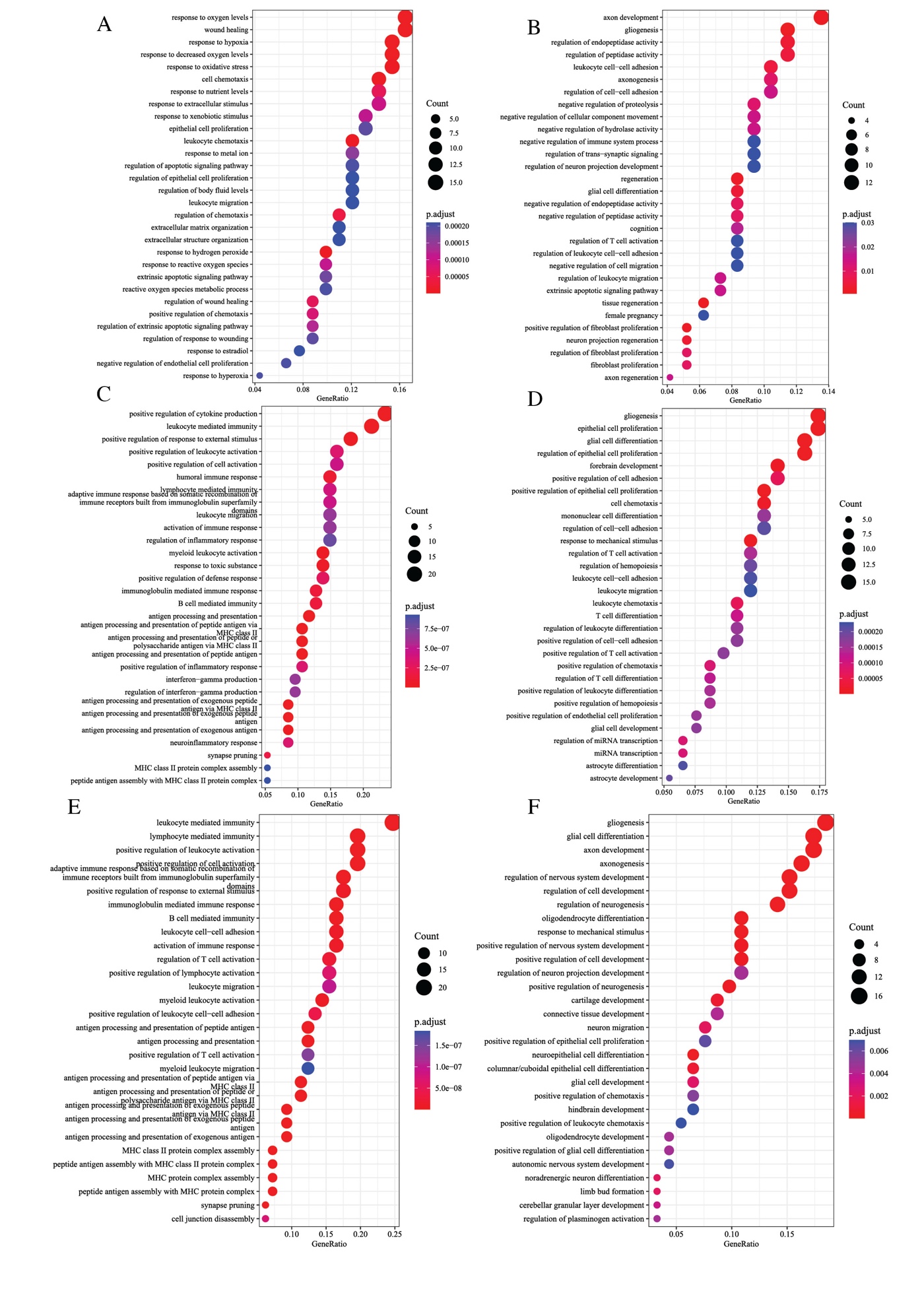

A. The figure displays the top 30 up-regulated BPs enriched by genes in the upper 5% quantile of the 12^th^ metagene.
B. The figure displays the top 30 down-regulated BPs enriched by genes in the lower 5% quantile of the 12^th^ metagene.
C. The figure displays the top 30 up-regulated BPs enriched by genes in the upper 5% quantile of the 13^th^ metagene.
D. The figure displays the top 30 down-regulated BPs enriched by genes in the lower 5% quantile of the 13^th^ metagene.
E. The figure displays the top 30 up-regulated BPs enriched by genes in the upper 5% quantile of the 15^th^ metagene.
F. The figure displays the top 30 down-regulated BPs enriched by genes in the lower 5% quantile of the 15^th^ metagene.
